# Structural variations contribute to subspeciation and yield heterosis in rice

**DOI:** 10.1101/2024.11.03.621459

**Authors:** Zhiwu Dan, Yunping Chen, Wenchao Huang

## Abstract

Yield heterosis has been extensively exploited in hybrid breeding, with inter-subspecies hybrids often exhibiting the most pronounced effects. However, developing elite hybrids remains a laborious and time-consuming process. The genetic basis of heterosis has been debated for over a century, hindered largely by the lack of high-quality genomes. Here, we assembled genomes for 12 representative *indica* and *japonica* rice accessions. Using sequence variants of the *Phr1* gene, we functionally validated two deletions responsible for phenol reaction variation between the subspecies. Comparative genomic analyses revealed extensive sequence variation among these inbred lines and highlighted the pivotal role of structural variants (SVs) in rice subspeciation. Importantly, the number of SVs between parental inbred lines significantly correlated with heterosis across 17 agronomic traits, with distinct correlation patterns for intra- and inter-subspecific F_1_ hybrids. We identified SVs associated with *S5-ORF5* and *OsBZR1* and validated their function to heterosis for seed setting rate and yield heterosis, respectively, underscoring the importance of SVs in breeding inter-subspecific hybrids. The genomic SVs altered gene expression and these transcriptional changes effectively explained the variances in heterosis. Furthermore, translocations outperformed other SVs and their heterozygous haplotypes exhibited heterosis over homozygous ones. Our findings establish SVs as pivotal drivers in subspeciation and highlight the overdominance model for harnessing rice heterosis.

## Introduction

Heterosis refers to the phenomenon where heterozygous F_1_ hybrids exhibit superior performance compared to their parental lines (Shull, 1948), significantly enhancing crop yield and contributing to global food security (Hochholdinger and Yu, 2025). Though several well-known genetic models—including dominance (complementation of deleterious loci) (Bruce, 1910; Jones, 1917; Gu et al., 2023; Wang et al., 2023), overdominance (heterozygosity of loci) (Shull, 1908, 1911; East, 1936; Semel et al., 2006; Krieger et al., 2010), and epistasis (interaction of loci) (Powers, 1944; Jiang et al., 2017)—have been proposed to explain heterosis, the genetic mechanisms underlying crop heterosis remain debated. Over a century ago, Shull proposed the overdominance model, emphasizing heterozygosity of “elements”, now always interpretated as genes or genomic sequences. He also suggested that the degree of heterosis correlates with the number of parental differential “characters”, now may be understood as genomic variants (Shull, 1911). Consistent with the opinion of Shull, East postulated that heterosis arises from the action of non-defective alleles and is linked to parental genetic disparity (East, 1909; East, 1936).

The advent of DNA marker technology in the late 20th century enabled investigations into the relationship between parental genetic disparity and heterosis in F_1_ hybrids using various molecular markers (Stuber et al., 1992; Xiao et al., 1995; Xiao et al., 1996; Zhang et al., 1996). Studies reported significant correlations between heterosis (or hybrid performance) and genetic distance or marker heterozygosity (Xiao et al., 1996; Zhang et al., 1996; Schrag et al., 2006; Schrag et al., 2009; Schrag et al., 2010; Dan et al., 2014; Liu et al., 2014). However, due to technology limitations, these studies relied on small number of marker sets, unable to reflect genome-wide sequence variation.

Advances in genome sequencing over the past two decades have dramatically improved the detection ability and completeness of genomic variants. High-density single-nucleotide polymorphism (SNP) and small insertion and deletion (InDel) markers (< 50 bp) have been widely used with various mathematical models to predict heterosis or hybrid performance in maize, rice, and wheat (Xu et al., 2014; Zhao et al., 2014; Zhao et al., 2015). Sequencing thousands of rice hybrids revealed that the number of superior alleles correlated strongly with grain number per panicle (Huang et al., 2015). In-depth sequencing analysis of 17 representative hybrid populations demonstrated that heterozygosity at heterosis-associated loci underlay rice heterosis (Huang et al., 2016). The sequencing of 1,143 *indica* inbred lines further underscored the importance of genomic variants in breeding superior hybrid rice (Lv et al., 2020). Based on sequencing results of 2,839 hybrids and their 9,839 segregation individuals, the authors found that the breeding of elite *indica*-*indica* hybrids was a process of pyramiding favorable alleles from broadened genetic resources (Gu et al., 2023). And they revealed widespread genomic complementarity in *indica*-*japonica* heterosis (Gu et al., 2023). Moreover, genomic sequencing analysis of two test-cross populations elucidated that rice heterosis was due to homozygous disadvantage at insufficient genetic background, rather than heterozygous advantage (Xie et al., 2022).

The contribution of large size genomic variants (≥ 50 bp), e.g., genome assembly-dependent structural variants (SVs), to rice heterosis remains poorly understood. Recent progress in sequencing technologies, including Pacific Biosciences (PacBio), Oxford Nanopore Technologies, and Hi-C platforms (Li and Durbin, 2024), has enabled the assembly of high-quality genomes. Over the past five years, the completeness and accuracy of genome assemblies have been significantly improved, leading to the release of multiple high-quality rice genomes (Qin et al., 2021; Shang et al., 2022; Shang et al., 2023; Sun et al., 2023). Given the important roles of SVs in tomato improvement and maize heterosis (Alonge et al., 2020; Wang et al., 2023), SVs are likely also critical for rice heterosis.

In this study, we selected 12 representative *indica* and *japonica* inbred lines and assembled their genomes for comparative analysis. We identified genomic variants, particularly SVs, and assessed their functional implications in rice sub-speciation. Using these inbred lines as parents, we constructed a diallel-cross population and analyzed the relationship between parental genomic variants and heterosis of 17 agronomic traits. We identified SVs in *S5-ORF5* and *OsBZR1* and validated their contributions to heterosis for seed setting rate and yield heterosis, respectively. In combination with gene expression data, we found that SVs drove transcriptional changes and that these effectively explained variance in rice heterosis. Furthermore, translocations outperformed other SVs and their heterozygous haplotypes conferred heterosis over the homozygous ones, supporting the overdominance model of heterosis in rice.

## Results

### Selection of 12 representative rice inbred lines

To investigate the genetic mechanisms underlying rice heterosis, we selected 12 representative *indica* and *japonica* accessions from 223 inbred lines (ILs) based on previous genotyping data (Dan et al., 2024) (Supplementary Fig. S1). These 12 ILs were equally clustered into *indica* and *japonica* groups (Fig. 1A), based on different types of genomic variants analyzed from the reported Illumina sequencing data (Dan et al., 2024). Among the selected ILs, seven—Yuetai B (YB), Mianhui725, Chenghui9348 (9311K), 610234, C418, Wuyunjing8, and Balilla—are well-established cultivars known for high grain yield or their widespread exploitation in breeding elite inbred or hybrid rice (https://www.ricedata.cn). Other ILs show highly recognizable phenotypes, e.g., large grain/primary branch/secondary branch number for Qianlijing, long panicle for R4115, and tall plant height for JR2 (Dan et al., 2024). These ILs represent phenotypically and geographically diverse varieties originating from China, India, and Italy (Fig. 1B).

**Figure 1.**
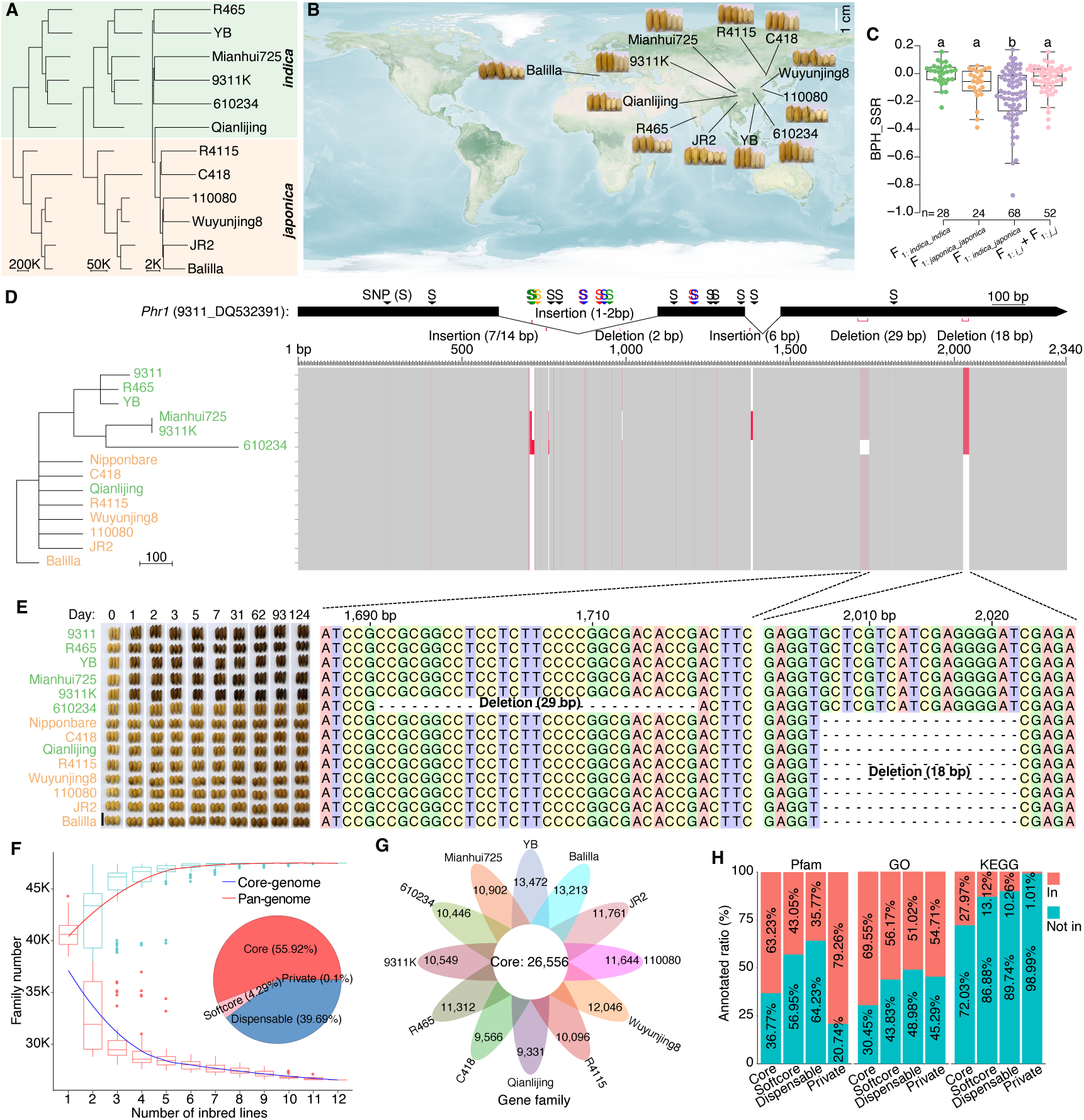
Genome assemblies and pan-genome of 12 rice inbred lines. **A)** Neighbour-joining trees of 12 representative rice inbred line (ILs) based on genomic SNPs (left side), InDels (middle), and CNVs (right side) analyzed from previously reported Illumina sequencing results. **B)** Geographic origins and seeds of the 12 ILs. **C)** Better-parent heterosis for seed setting rate (BPH_SSR) of intra- and inter-subspecific F_1_ hybrids. The pooled average values of *indica*-*indica* (*i_i*) and *japonica*-*japonica* (*j_j*) F_1_ hybrids were also included for comparison. Analysis of variance (ANOVA) with LSD in post-hoc test (*P*-value < 0.05) was performed. The number of F_1_ hybrids are indicated. **D)** Gene structures and neighbour-joining tree of the 12 ILs based on gene sequences of *Phr1*. Nipponbare (*japonica*) and 9311 (*indica*) served as reference accessions. Red regions denote genomic variants, and dashed lines represent deletions. **E)** Phenol reaction phenotypes of 14 ILs. The bar represents 1 cm. **F)** Variation of gene family sizes in the pan- and core-genome. **G)** Core and non-core gene families across the 12 ILs. **H)** Gene annotation ratios cross three databases. Three databases, including Pfam, GO, and KEGG, were selected for annotation analysis.

Using a complete diallel-cross design, we generated F_1_ hybrids from the 12 ILs and calculated better-parent heterosis (BPH) of 17 agronomic traits for the finally obtained 120 F_1_ hybrids. We observed that F_1_ hybrids derived from inter-subspecies exhibited significantly reduced seed setting rate and increased plant height compared to those of intra-subspecific hybrids (Fig. 1C and Supplementary Fig. S2), which is consistent with known breeding challenges. Furthermore, heterosis of the inter-subspecific hybrids showed significant differences from the pooled averages of *indica-indica* and *japonica-japonica* hybrids.

### Genome assembly and annotation of 12 inbred lines

To generate genome assemblies, we obtained genomic data using single-molecule real-time (SMRT) technology on the PacBio Revio platform for the 12 ILs, achieving an average sequencing depth at 30.46× (Supplementary Tables S1-S2). We performed *de novo* assembly for two representative ILs, YB (*indica*) and Balilla (*japonica*), with the assistance of Hi-C data, yielding an average sequencing depth at 127.80× (Supplementary Fig. S3 and Supplementary Table S3). The genomes at the chromosome level of the remaining ten ILs were assembled using YB and Balilla as reference genomes for *indica* and *japonica* subgroups, respectively. The average contig N50, scaffold N50, and total assembled genome sizes for final genome assemblies of the 12 ILs were 28.74 Mb, 31.75 Mb, and 396.36 Mb, respectively, with an average anchor ratio of 98.18%. The high-quality and completeness of these genome assemblies were further confirmed by metrics such as average quality value of bases (63.06), re-mapping rate of HiFi reads (99.51%), Benchmarking Universal Single-Copy Orthologs (BUSCO) (98.61%), and long terminal repeats (LTRs) assembly index (22.13) (Supplementary Fig. S4).

We annotated repetitive sequences (tandem repeats and transposable elements), protein-coding genes, and non-coding RNAs for YB and Balilla (Supplementary Table S4). Repetitive sequences constituted 57.35% (YB) and 55.44% (Balilla) of their genomes, with transposable elements (TEs) comprising 55.09% and 53.10%, respectively. Among these TEs, DNA transposons and LTRs were the predominant types, accounting for 24.26-25.67% and 24.9-28.64% of the genomes, respectively. Three strategies, including homology-based (using five previously reported genomes), *de novo*, and transcriptome-based (full-length RNA sequencing) predictions, were integrated for the annotation of protein-coding genes. This resulted in YB containing 44,332 genes and Balilla containing 43,706 genes. We revealed that 94.4% of the annotated genes in YB and 95.35% in Balilla have assigned functions in one of the following six databases: InterPro, Gene Ontology (GO), Kyoto Encyclopedia of Genes and Genomes (KEGG), Swiss-Prot, TrEMBL, and Non-Redundant. Compared to the five established genomes (Nipponbare, 93-11, Minghui63, Zhenshan97, and R498), these newly assemblies showed significant different in the lengths of genes and coding sequences, highlighting their value to current rice genomic resources (Supplementary Fig. S5). Moreover, miRNA and rRNA were the two predominant types of non-coding RNAs in the genome assemblies of both YB and Balilla, which accounted for 0.26-0.28% and 0.34-0.96% of the genomes, respectively.

### Accuracy validation of genome assemblies using *Phr1*

To further validate assembly accuracy, we analyzed *Phr1*, a gene underlying phenol reaction phenotype between *indica* and *japonica* (Yu et al., 2008). Phylogenetic result of *Phr1* sequences from the 12 ILs, along with 9311 and Nipponbare, displayed slightly changed clustering patterns in subspecies, with Qianlijing grouped with *japonica* (Fig. 1D). Using the sequence of 9311 as reference, *Phr1* contained 24 SNPs and six InDels. Among these InDels, there existed three insertions (1-2, 6, and 7 or 14 bp) and three deletions (2, 18, and 29 bp), with only the 18 and 29 bp deletions located in exons. The result of phenol reaction experiment revealed that seed hulls of five ILs without the 18/29 bp deletions immediately turned black after one day’s treatment, while the other nine remained yellow or brown even after 124 days (Fig. 1E). These results showed the high assembly accuracy of these new genomes and confirmed important roles of the 18/29 bp deletions in outcomes of phenol reaction.

### Pan-genome assembly of 12 inbred lines

A pan-genome of the 12 ILs was constructed, comprising 47,490 gene families (Fig. 1F). Among the whole gene families, 26,556 (with 345,960 genes) were identified in all 12 ILs and defined as core-gene families (55.92% of gene families and 70.70% of genes), 2,037 (with 24,040 genes) identified in 11 ILs were defined as softcore gene families (4.29% of gene families and 4.91% of genes), 18,850 (with 118,436 genes) identified in 2-10 ILs were defined as dispensable families (39.69% of gene families and 24.20% of genes), and 47 (with 892 genes) identified in only one IL were defined as private gene families (0.10% of gene families and 0.18% of genes). The number of gene family increased with the addition of genomes, reaching a plateau at *n* = 10, indicating the representativeness of the selected ILs (Fig. 1F). Unlike the trend observed in pan genome, the number of core gene family decreased as the number increased, with the lowest point reached at *n* = 12, underscoring the specificity of each IL (Fig. 1F). For the number of non-core gene families among these ILs, YB and Qianlijing had the highest and lowest amount, respectively, showing functional variations between ILs (Fig. 1G). Furthermore, we analyzed the proportion of genes with annotated functions in protein domain annotation (Pfam), GO, and KEGG. The annotation ratios decreased progressively from core to softcore to dispensable genes (Pfam: 63.23%, 43.05%, and 35.77%; GO: 69.55%, 56.17%, and 51.02%; and KEGG: 27.97%, 13.12%, and 10.26%), demonstrating the conservation of core genes amidst diversification (Fig. 1H).

### Pan-structural variants in the 12 inbred lines

Genomic variants—especially SVs (presence/absence variants, copy number variants, translocations, and inversions)—were identified through comparing the reference genome YB to the other 11 ILs. Consistent with the patterns observed for SNPs and InDels (Fig. 2A), both the number and size (about 57-168 Mb) of SVs gradually increased from *indica* to *japonica* (Fig. 2B-C). All *japonica* ILs harbored more SVs than the *indica*. Notably, two large inversions were detected on chromosome 6 (inversion_6: 5.83 Mb) and 8 (inversion_8: 5.90 Mb) (Fig. 2D), which were between the six *indica* and six *japonica* or exclusive to JR2 (Supplementary Fig. S6-S10), respectively, indicating that inversions contribute to the evolutionary divergence of *indica* and *japonica*.

**Figure 2.**
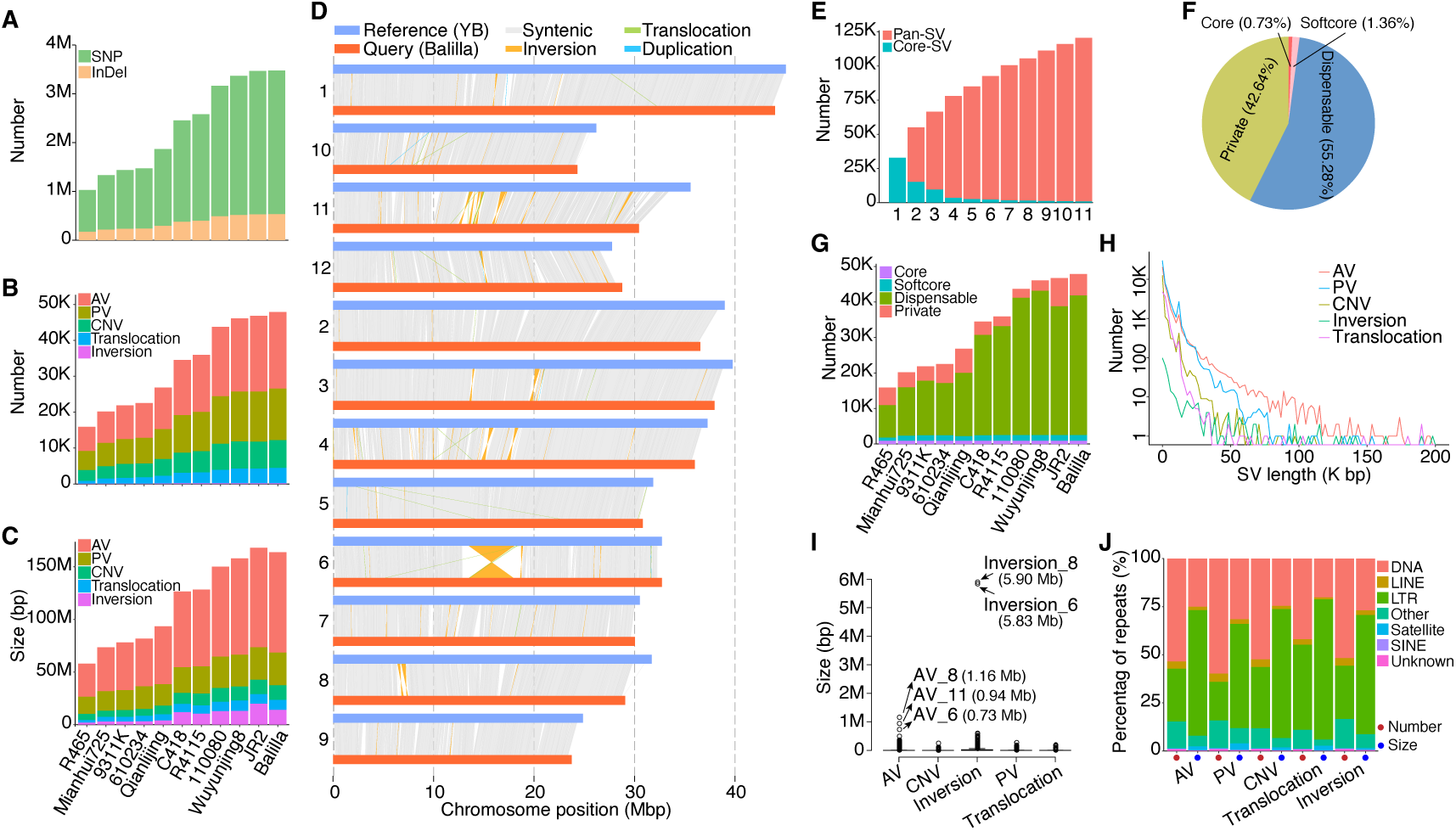
Genomic variants of 12 inbred lines. **A-C)** Number and size distribution of genomic variants in 11 inbred lines with genome assembly of YB as the reference. **D)** Comparison of genomic profiles between YB and Balilla. **E)** Variation of SVs in the pan- and core-SVs with varied number of inbred lines. **F)** Percentages of core, softcore, private, and dispensable SVs. **G)** Number of core, softcore, private, and dispensable SVs in 11 inbred lines. **H-I)** Number and size of SVs. **J)** Percentages of different types of repeats in SVs. The number and size are analyzed.

Unlike the changing trend of pan-genome, the number of pan-SVs did not plateau even at *n* = 11, underscoring the necessity of including more accessions for a comprehensive exploration of SVs (Fig. 2E). Core, softcore, dispensable, and private SVs were assigned using the same criteria as the pan-genome. Among the total SVs, 878 (0.73%) identified in 11 ILs were classified as core SVs, while 1,640 (1.36%) detected in 10 ILs were categorized as softcore SVs (Fig. 2F). Additionally, 66,655 (55.28%) SVs identified in 2-9 ILs were classified as dispensable SVs, and 51,413 (42.64%) identified in only one IL were considered private SVs. Different from the trends of core, softcore, and private SVs, the number of dispensable SVs increased from *indica* to *japonica* (Fig. 2G), suggesting the roles of these dispensable SVs played in rice subspeciation. The length of different SV types varied (Fig. 2H), with three large absence variants (AVs) detected on chromosome 6 (AV_6, 0.73 Mb), 8 (AV_8, 1.16 Mb), and 11 (AV_11 and 0.94 Mb; Fig. 2I). Given that repetitive sequences accounted for more than 50% of the genomes, we analyzed the distributions of SVs and found that these SVs were predominantly detected in DNA transposons and LTRs (Fig. 2J), suggesting regulatory functions of SVs in non-coding regions of the genomes.

### Associations between structural variants and heterosis of 17 agronomic traits

We performed paired genomic comparisons and identified both unique and common genomic variants for each pair of ILs, representing different and shared variants, respectively (Fig. 3A and Supplementary Table S5). On average, unique genomic variants between any two ILs comprised 2,018,539 SNPs, 391,298 InDels, and 40,082 SVs (17,690 AVs, 11,975 PVs, 6,327 CNVs, 3,941 translocations, and 149 inversions). The number of these unique variants was highly correlated (Fig. 3B), with the exception of translocation, inferring co-locations of these variants (Supplementary Fig. S11). Approximately half of these variants located in intergenic regions (57.45% of SNPs, 50.42% of InDels, and 50.56% of SVs), emphasizing the significance of these regions in differentiating the ILs. In contrast to other SV types, the majority (77.94%) of unique inversions occurred in exonic regions. Furthermore, genetic relatedness of the 12 ILs assessed using unique SVs, particularly translocations, showed variation compared to those based on unique SNPs or InDels, indicating the distinct functional contributions of different variant types to genomic divergence (Fig. 3C and Supplementary Fig. S12).

**Figure 3.**
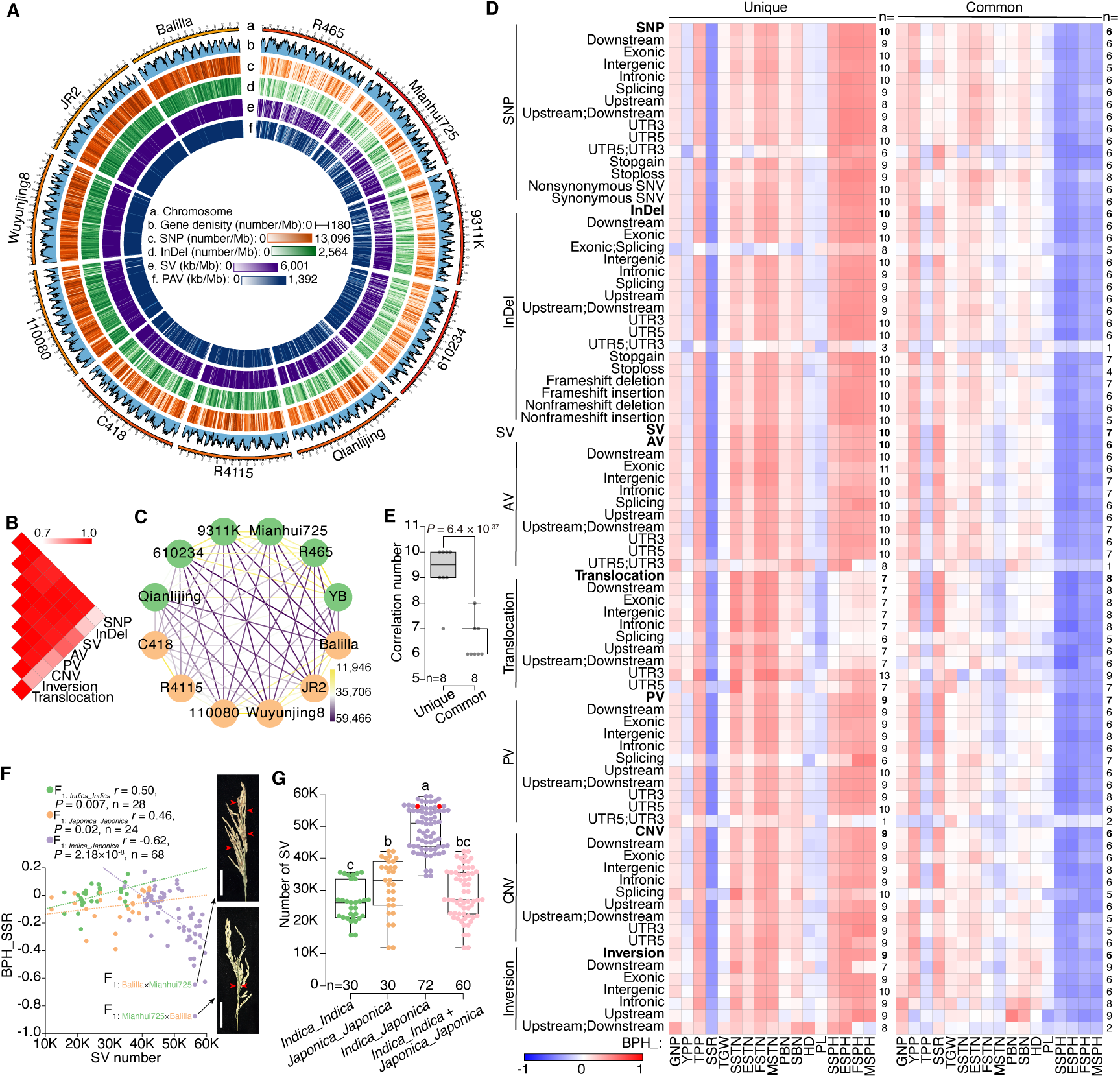
Correlations between parental genomic variants and heterosis of 17 agronomic traits. **A)** Genomic comparisons of YB against the other 11 inbred lines (ILs). **B)** Heatmap for the correlations between different types of unique genomic variants. **C)** Genetic relatedness of 12 ILs based on unique SVs. **D)** Correlation patterns between the number of parental genomic variants and heterosis of 17 traits. Parental unique and common variants were classified into different types according to their location or function. Spearman correlation analysis was performed with better-parent heterosis (BPH) of the following traits: grain number per panicle (GNP), tiller number per plant (TPP), seed setting rate (SSR), thousand-grain weight (TGW), seedling stage tiller number (SSTN), elongation stage tiller number (ESTN), flowering stage tiller number (FSTN), maturation stage tiller number (MSTN), yield per plant (YPP), seedling stage plant height (SSPH), elongation stage plant height (ESPH), flowering stage plant height (FSPH), maturation stage plant height (MSPH), panicle length (PL), primary branch number (PBN), secondary branch number (SBN), and heading date (HD). The number besides each row indicates the number of significant correlations (*P*-value < 0.05). **E)** Comparison of correlation frequencies between unique and common genomic variants. *P*-value indicates independent *t*-test (weighted). **F)** Correlations between the number of unique SVs and heterosis for seed setting rate in intra- and inter-subspecific F_1_ hybrids. A pair of reciprocal hybrids (F_1: Mianhui725×Balilla_ and F_1: Balilla×Mianhui725_) with extremely low heterosis are shown. Mature seeds with black and white hulls are full and empty seeds, respectively. *P*-values indicate Spearman correlation. **G)** The number of unique SVs in intra- and inter-subspecific F_1_ hybrids. The two reciprocal hybrids are marked in red. ANOVA with LSD in post-hoc test (*P*-value < 0.05) was performed.

Correlation analysis between the number of parental genomic variants and heterosis of the 17 traits revealed divergent patterns for unique versus common variants (Fig. 3D). According to the number of significant correlations, the unique variants exhibited stronger correlations with heterosis than common variants (Fig. 3E). Meanwhile, no notable change in correlation patterns was observed between stop-gain and stop-loss variants in SNPs and InDels, synonymous and nonsynonymous SNPs, or frameshift and nonframeshift InDels. These findings emphasized the importance of genomic differences between the two parents in determining heterosis.

Though unique SVs accounted for only 1.99% and 10.24% of the total SNPs and InDels, respectively, their correlations with heterosis were comparable, probably due to their larger size. Among the five types of SVs, inversions, with fewer number than translocations, showed stronger correlations with heterosis for plant height, inferring a size-dependent influence of genomic variants on heterosis of vegetative traits. Different from correlation patterns of heterosis for other yield-related traits, the number of unique variants showed negative correlations with heterosis for seed setting rate, showing reproductive isolation due to genomic disparity. Notably, we observed that an increase in unique SVs corresponded with a slight rise in heterosis for seed setting rate of intra-subspecific F_1_ hybrids, but a dramatic decrease in *indica*-*japonica* F_1_ hybrids (Fig. 3F). The inter-subspecific hybrids possessed significantly more unique SVs in their parents than their intra-subspecific counterparts (Fig. 3G and Supplementary Fig. S13), suggesting the roles of these SVs in heterosis of inter-subspecies. The unique SVs between the parental lines also exhibited stronger correlations with heterosis for tiller number in inter-subspecific F_1_ hybrids, showing the potential for enhancing grain yield through utilizing heterosis for tiller number (Supplementary Fig. S14). However, the number of unique SVs displayed weak or no correlation with yield heterosis in inter-subspecific F_1_ hybrids, likely due to reproductive isolation-driven reductions in seed setting rate.

### Absence variant of *S5-ORF5* underlies heterosis for seed setting rate

Given the established role of the *S5* loci in embryo-sac fertility and seed setting rate of inter-subspecific F_1_ hybrids (Chen et al., 2008; Yang et al., 2012), we performed phylogenetic analyses of its three genes (*S5-ORF3*, *S5-ORF4*, and *S5-ORF5*) in the 12 ILs (Fig. 4A). Unexpectedly, two ILs—YB and Qianlijing—clustered together and were identified as wide compatibility varieties (WCVs). With gene sequence of Balilla as the reference, we detected eight SNPs, three deletions (1, 10, and 42 bp), and one 94 bp AV in *S5-ORF5* (Fig. 4A), which acts as the “killer” gene. The 94 bp AV was exclusive to the two WCVs, potentially rescuing embryo-sac viability in their inter-subspecific F_1_ hybrids.

**Figure 4.**
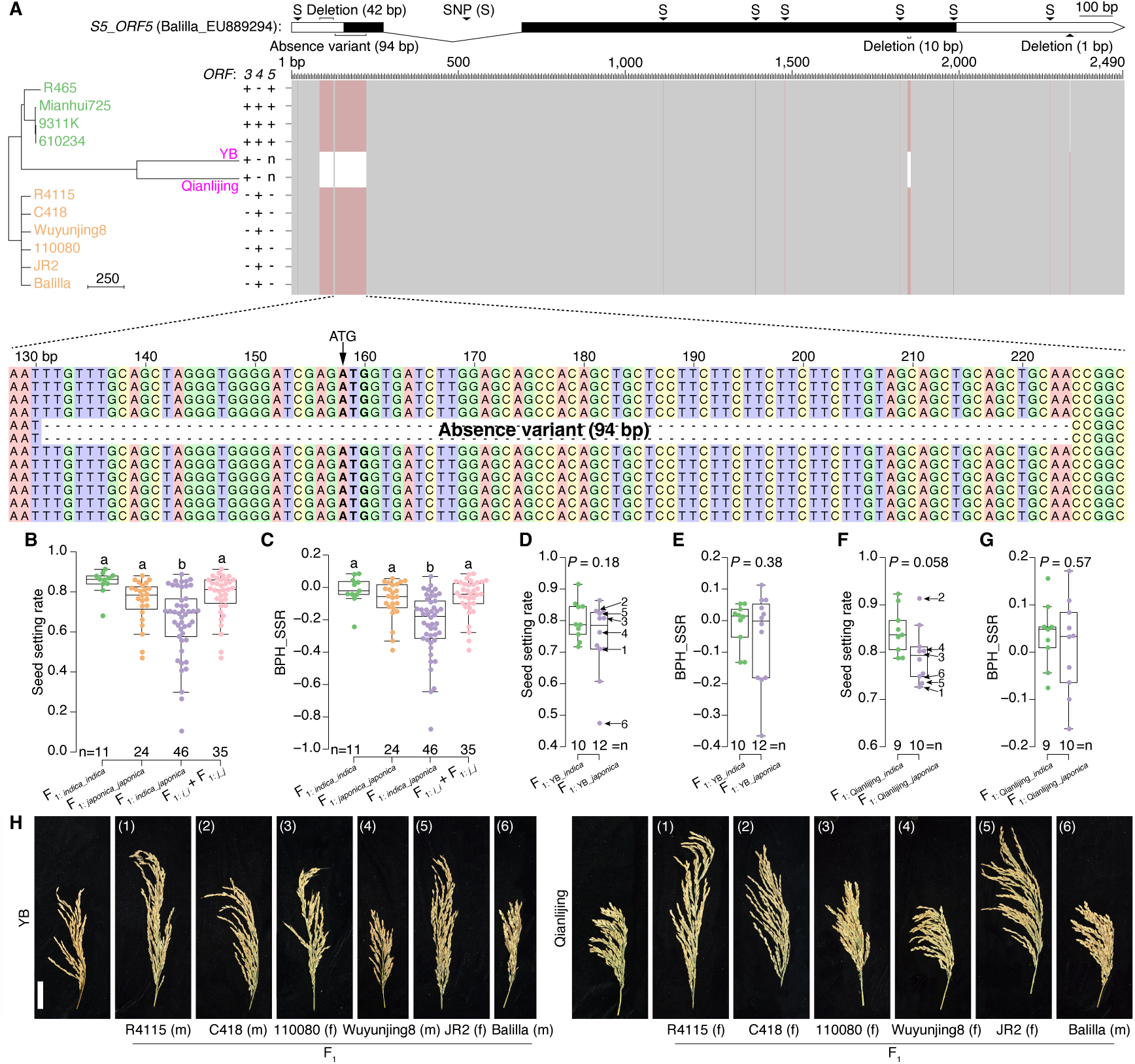
Contribution of AVs in *S5-ORF5* to heterosis for seed setting rate. **A)** Neighbour-joining tree of 12 inbred lines (ILs) based on *S5* loci (three genes) and gene structure of *S5-ORF5*. Genotypes of *S5-ORF3*, *S5-ORF4*, *S5-ORF5* for 12 ILs were assigned with previously reported rules. The “n” represents wide compatibility varieties. The sequence of Balilla serves as the reference and sequence variants are indicated in red. **B-C)** Comparisons of seed setting rate and better-parent heterosis for seed setting rate in intra- and inter-subspecific F_1_ hybrids. Two wide compatibility varieties, namely YB and Qianlijing, were excluded for analysis. ANOVA with LSD in post-hoc test (*P*-value < 0.05) was performed. **D-E)** Comparisons of seed setting rate and better-parent heterosis for seed setting rate between intra- and inter-subspecific F_1_ hybrids with YB as a parent. **F-G)** Comparisons of seed setting rate and heterosis for seed setting rate between intra- and inter-subspecific F_1_ hybrids with Qianlijing as a parent. *P* values in **D**-**G** indicate independent sample *t*-test. **H)** Panicles of inter-subspecific F_1_ hybrids with YB or Qianlijing as a parent. The letter f and m represent female and male parent, respectively. Seed setting rate of these F_1_ hybrids are displayed in **D** and **F**.

To investigate contribution of the 94 bp AV to heterosis, we first compared seed setting rate between intra- and inter-subspecific F_1_ hybrids lacking the WCV parentage. The seed setting rate of *indica*-*indica* and *japonica*-*japonica* F_1_ hybrids were similar, both significantly higher than those of *indica*-*japonica* F_1_ hybrids (Fig. 4B). The heterosis for seed setting rate was also lower in inter-subspecific F_1_ hybrids (Fig. 4C), compared to those of intra-subspecific F_1_ hybrids and their pooled average values. However, when YB or Qianlijing was used as a parent, the *indica*-*japonica* F_1_ hybrids achieved seed setting rate as high as the *indica*-*indica* F_1_ hybrids (Fig. 4D-H). Importantly, heterosis for seed setting rate was also comparable between the inter- and intra-subspecific F_1_ hybrids. We therefore identified YB and Qianlijing as two *indica*-type WCVs and confirmed important role of the 94 bp AV in heterosis for seed setting rate.

### Presence and absence variants of *OsBZR1* contribute to yield heterosis

*OsBZR1*, a positive regulator of the brassinosteroid signaling pathway (He et al., 2002), regulates phenotypic differentiation between *indica* and *japonica* subspecies (Yang et al., 2025), including yield per plant, tiller number, grain number, thousand-grain weight, panicle length, and plant height. Clustering of the 12 ILs based on phenotypic data of these traits supported *OsBZR1*’s role in subspecific differentiation (Supplementary Fig. S15). Given that sequence variants in the *OsBZR1* promoter region—particularly a 4,808 bp retrotransposon—are critical for its function, we extracted and compared *OsBZR1* sequences in the 12 ILs. Phylogenetic analysis revealed that 11 of the 12 ILs clustered into distinct *indica* and *japonica* groups, with JR2 as an exception (Fig. 5A). We thus classified the ILs into three haplotypes: haplotype_i (*indica*), haplotype_j (*japonica*), and haplotype_JR2 (divergent).

**Figure 5.**
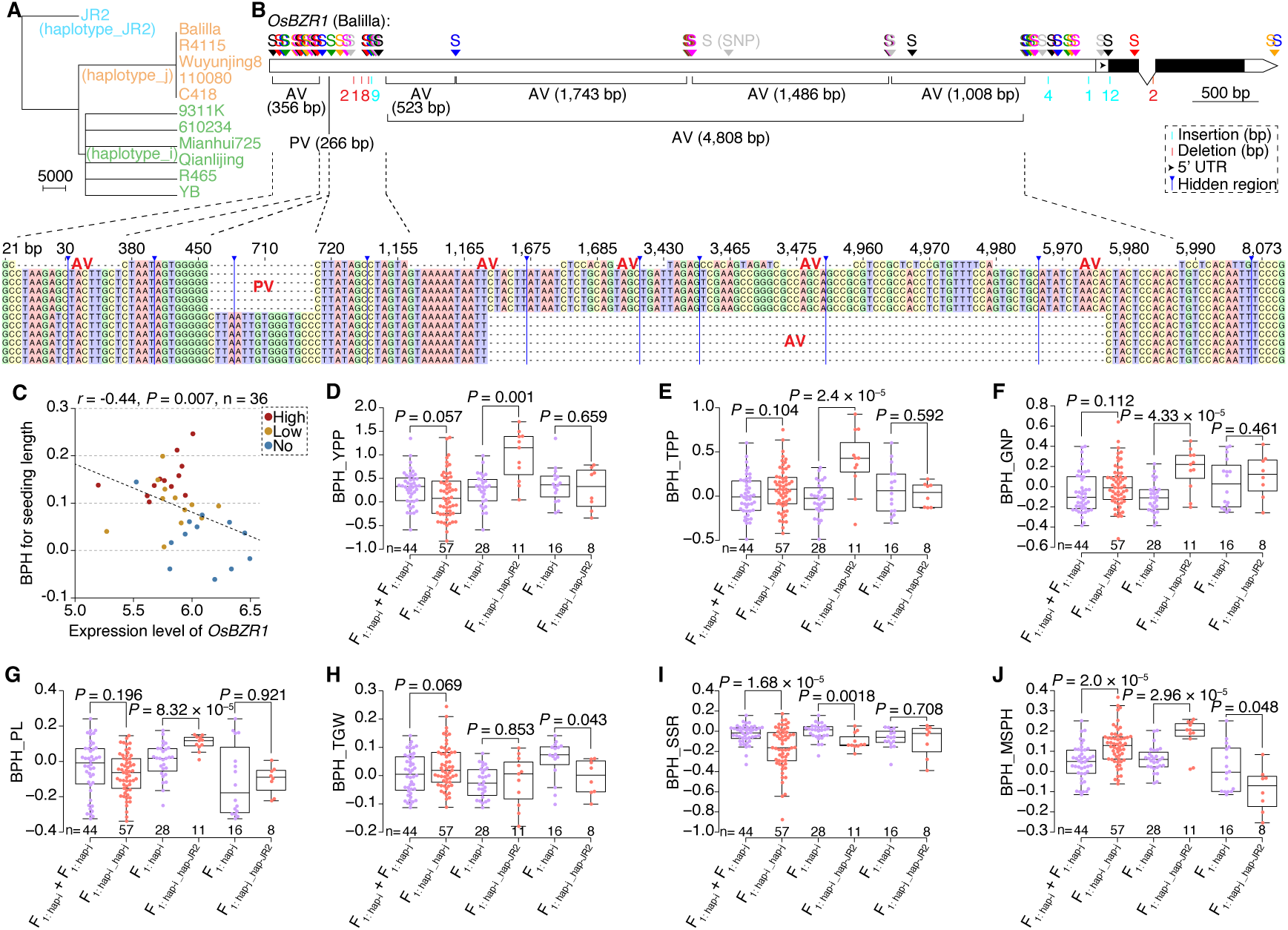
Contributions of SVs in *OsBZR1* to yield heterosis. **A)** Neighbour-joining tree of 12 inbred lines (ILs) based on sequences *OsBZR1*. The 12 ILs were divided into three haplotypes. **B)** Structure variations of *OsBZR1* across 12 ILs. The sequence of Balilla serves as the reference. Single nucleotide polymorphism (SNP), insertion and deletion (InDel), absence variant (AV), and presence variant (PV) were indicated. **C)** Correlation between expression level of *OsBZR1* and better-parent heterosis (BPH) for seedling length. Six pairs of reciprocal hybrids, which were classified as with high-, low-, and no-BPH for seedling length, were adopted for correlation analysis. *P* value indicates Spearman correlation. **D-J)** Comparisons of BPH for seven *OsBZR1*-associated traits between heterozygous and homozygous haplotypes. The seven traits are yield per plant (YPP), tiller number per plant (TPP), grain number per panicle (GNP), thousand-grain weight (TGW), panicle length (PL), maturation stage plant height (MSPH), and seed setting rate (SSR). Hap, i, and j represent haplotype, *indica*, and *japonica*, respectively. *P* values in **D**-**J** are for independent sample *t*-test.

Using the sequence of Balilla as reference, we identified 82 SNPs, four insertions (1, 4, 9, and 12 bp), four deletions (1, 2, 2, and 8 bp), one 266 bp PV, and six AVs (356, 523, 1,008, 1,486, 1,743, and 4,808 bp) (Fig. 5B). The 266 bp PV was exclusive to the six *japonica* accessions, while the 4,808 bp AV was present in the six *indica* lines, indicating contributions of these SVs to subspeciation. The remaining five AVs, with four of which mainly derived from the 4,808 bp AV, were unique to JR2.

Since the retrotransposon influences gene transcription, we analyzed the relationship between expression levels of *OsBZR1* and heterosis for seedling length in six pairs of reciprocal F_1_ hybrids, which were divided into high-, low-, and no-heterosis groups (Dan et al., 2024). We observed a significantly negative correlation, suggesting contributions of low expression levels of *OsBZR1* to heterosis for seedling length (Fig. 5C). When expression levels were compared between F_1_ hybrids and corresponding parents, we detected overdominant effect, dominant effect, and unclassifiable effects, indicating diverse inheritance patterns of *OsBZR1* across different hybrid backgrounds (Supplementary Fig. S16).

To assess the impact of genomic variants in *OsBZR1* on heterosis of field traits, we compared heterosis of the seven traits between F_1_ hybrids derived from ILs carrying different haplotypes (Fig. 5D-J). Except for seed setting rate and plant height, heterosis of haplotype_i/haplotype_j heterozygous F_1_ hybrids did not significantly differ from the average of their homozygous counterparts (haplotype_i and haplotype_j). Strikingly, F_1_ hybrids with the haplotype_i/haplotype_JR2 exhibited significantly stronger heterosis than the haplotype_i homozygotes for most traits, including yield, tiller number, and plant height. In contrast, hybrids combining haplotype_j and haplotype_JR2 showed no significant yield heterosis advantage over haplotype_j homozygotes, highlighting the importance of the haplotype_i/haplotype_JR2 heterozygous state for robust yield heterosis.

### Changes in gene expression by structural variants explain variance in heterosis

To further explore the impacts of SVs on gene expression, we performed RNA-sequencing and differential expression analyses on seedling samples of the 12 ILs (Fig. 6A). The differentially expressed genes (DEGs) clustered the ILs into two major groups, with C418 and R4115 in *indica* group, inferring changes in genetic information from the genomic to transcriptomic levels (Supplementary Fig. S17). Drawing on previously reported inheritance patterns of metabolites in rice F_1_ hybrids (Dan et al., 2024), we demonstrated that transcriptomic additive and partially dominant effects played more important roles than dominant effects in heterosis across the 17 traits, reinforcing contributions of the two main effects to heterosis across different genetic levels (Fig. 6B and Supplementary Fig. S17).

**Figure 6.**
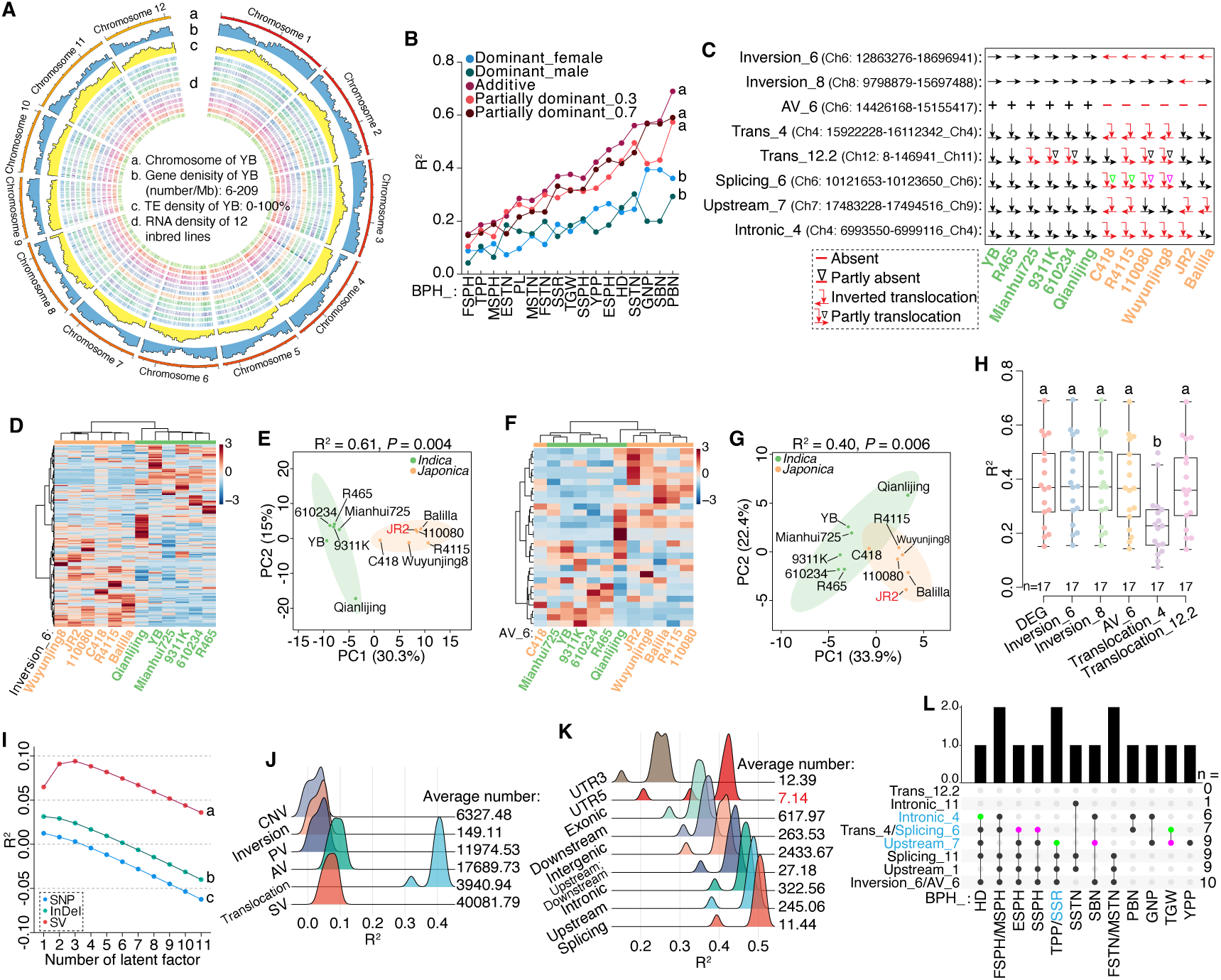
Relationships among genomic SVs, gene expression, and heterosis. **A)** Transcriptomic profiles of 12 inbred lines (ILs). Gene expression data were obtained from 15-day-old seedlings. **B)** Changes in R^2^ (adjusted) for heterosis of 17 traits. Different inheritance patterns, including dominant (female and male), additive, and partially dominant (0.3 and 0.7) effects, of expressed genes were used as predictive variables in partial least square (PLS) analysis on better-parent heterosis (BPH) of 17 agronomic traits. The best regression model (with highest adjusted R^2^) was selected as the final predictive model for heterosis of each trait. **C)** Genomic structures of two inversions, one absence variant (AV), and five translocations (trans) across 12 ILs. **D-G)** Heatmaps and PCA plots of 12 ILs based on expression levels of genes within inversion_6 and AV_6. *P*-values indicate PERMANOVA. **H)** Comparisons of R^2^ (adjusted) for predicting BPH of 17 traits with differentially expressed genes (DEGs) of 12 ILs, genes in inversion_6, genes in inversion_8, genes in AV_6, genes in translocation_4, and genes in translocation_12.2. **I)** Changes in R^2^ (adjusted) of PLS models with varied number of latent factors. The additive expression levels of DEGs from 12 ILs were used as variables to predict the number of parental genomic variants. The genomic variants include single nucleotide polymorphism (SNP), insertion and deletion (InDel), and structural variant (SV). **J)** R^2^ (adjusted) for different types of SVs. **K)** R^2^ (adjusted) for different types of translocations. **L)** Associations between SVs and heterosis. Genes linked to intronic_4, splicing_6, and upstream_7 were used to perform Spearman correlation analysis with heterosis. Their positive and negative correlations with heterosis are indicated in purple and green, respectively. Analysis of variance (ANOVA) with LSD in post-hoc test (*P*-value < 0.05) was performed in **B**, **H**, and **I**. Abbreviations: yield per plant = YPP, secondary branch number = SBN, grain number per panicle = GNP, tiller number per plant = TPP, seed setting rate = SSR, heading date = HD, thousand grain weight = TGW, seedling stage tiller number = SSTN, elongation stage tiller number = ESTN, flowering stage tiller number = FSTN, maturation stage tiller number = MSTN, seedling stage plant height = SSPH, elongation stage plant height = ESPH, flowering stage plant height = FSPH, maturation stage plant height = MSPH, panicle length = PL, and primary branch number = PBN.

We next focused on expression levels of genes within the two large inversions: inversion_6 (304 genes) and inversion_8 (331 genes) (Fig. 6C). Consistent with the presence/absence pattern of inversion_6, the 12 ILs were accurately classified into *indica* and *japonica* groups based on expression levels of these genes (Fig. 6D-E). Unexpectedly, genes within inversion_6 with additive or partially dominant effects were predictive of heterosis in partial least square (PLS) models (Supplementary Fig. S18). Based on genes in inversion_8, JR2 was found to be closer to *indica* than to *japonica* (Supplementary Fig. S19). Though only a limited number of genes differed in expression levels between the *indica* and *japonica*, the genes within inversion_8 effectively explained the variance in heterosis (Supplementary Fig. S19). Among the three large size AVs, genes in AV_8 (22 genes) and AV_11 (22 genes) were seldomly expressed in seedlings. For AV_6 (34 genes), most genes were expressed and effectively divided these ILs into two groups, with C418 in *indica* group (Fig. 6F-G). The genes in AV_6 also demonstrated predictability for heterosis comparable to the genome-wide DEGs of 12 ILs and the two inversions (Fig. 6H and Supplementary Fig. S20). These results confirmed the direct impact of SVs on gene expression, which in turn explained the variance in heterosis.

### Translocation outperforms other SVs in heterosis

To comprehensively investigate associations between different types of parental genomic variants and expression changes in F_1_ hybrids, we performed PLS analysis between additive expression levels of 1,759 DEGs from the 12 ILs and the number of genomic variants in each pair of ILs. Based on the R^2^ (adjusted) values of PLS models with varying number of latent factors, the predictability of these DEGs was higher for SVs than those for SNPs and InDels (Fig. 6I). When SNPs and InDels were further subdivided by location or function, the predictability remained low, except for variants located in multiple regions with their number below ten (Supplementary Fig. S21-22). In contrast, the predictability varied considerably among different types of SVs, with translocation predicted at the highest accuracy (Fig. 6J). Among the top five largest translocations, genes associated with translocation_8.1, translocation_12.1, and translocation_8.2 were rarely expressed in seedling samples (Supplementary Fig. S23). Translocation_4 was an inverted translocation on chromosome 4, detected between six *indica* and four *japonica*, except for JR2 and Balilla (Fig. 6C). This translocation contained 17 genes, eight of which had detectable expression levels (Supplementary Fig. S24). Due to loss of this translocation, Balilla was clustered with *indica* while JR2 was remained in *japonica* group, suggesting divergent effect of the same translocation on gene expression under different genetic background. The predictability of genes in translocation_4 for heterosis was lower than those of DEGs, the two inversions, and the AV_6 (Fig. 6H), which was mainly caused by the limited number of expressed genes. Translocation_12.2, between chromosomes 12 and 11, was present fully in some ILs and partially in others. Based on the normalized expression levels of its 17 genes, 110080 and Wuyunjing8 with partial translocations were clustered with *indica* ILs, while three *indica* with translocations were remained in their group (Supplementary Fig. S24), inferring differed translocation effect on gene expression between subspecies. Despite its smaller size, the predictability of expressed genes in translocation_12.2 for heterosis was as effectively as those within inversions and AVs (Fig. 6H).

When different types of SVs (with average variant number over ten) were detailly analyzed by location, the predictability was higher for exonic region of AV (Supplementary Fig. S25), UTR5 region of CNV (Supplementary Fig. S26), splicing region of PV (Supplementary Fig. S27), and intergenic region of inversion (Supplementary Fig. S28), suggesting diversified function of SVs on gene expression through different locations. The predictability for translocation also varied across locations, with splicing, upstream, and intronic regions having higher accuracy (Fig. 6K). To further investigate functions of different types of translocations, we selected three top size ones in combination with the identified DEGs: splicing_6, upstream_7, and intronic_4 (Fig. 6C), which had available gene expression levels. These three representative translocations were invariant in *indica* but variable in *japonica*. Using a haplotype analysis strategy similar to that for *OsBZR1*, we classified the 12 ILs into two haplotypes based on the presence/absence of each translocation. Then, we compared heterosis of the 17 agronomic traits between hybrids with heterozygous and homozygous haplotypes to investigate their relationships with heterosis. The associations varied across translocations, with the associated genes showing either positive or negative correlations with heterosis (Fig. 6L). We also analyzed three other representative translocations, including splicing_11, upstream_1, and intronic_11, which were detected in at least five ILs but with nearly undetectable gene expression (Supplementary Fig. S23). These translocations showed distinct associations with heterosis (Fig. 6L). After the haplotype analysis was extended to all the above-mentioned typical SVs, we found the largest number of associations for inversion_6 and AV_6 (n = 10; Fig. 6L). Then, the upstream_7, splicing_11, and upstream_1 were associated with heterosis of nine traits. The upstream_7 was the only translocation for yield heterosis, warranting further validation of this region or its associated gene by sequence modulation technologies. Taken together, the heterozygous haplotypes of SVs displayed significant heterosis to their homozygous ones, supporting the overdominance model in rice heterosis.

## Discussion

The genetic mechanism of heterosis has been a central debate in plant genetics for over a century. The overdominance model emphasizes the contribution of genomic heterozygosity and the co-existence of non-defective alleles (Shull, 1908; East, 1909; Shull, 1911; East, 1936). In this study, genomic variants in parental lines are at heterozygous states in F_1_ hybrids. SNPs, InDels, PAVs, CNVs, inversions, and translocations represent heterozygosity of single bases (1 bp), several bases (2-50 bp), some bases (> 50 bp), sequence repeat number, sequence direction, and sequence location, respectively (Supplementary Fig. S29). When genomic variants are combined for haplotype analysis, F_1_ hybrids with heterozygous haplotypes show heterosis to those with homozygous haplotypes, underscoring the critical role of genomic heterozygosity in rice heterosis. For the *S5-ORF5* locus, typical *indica*-*japonica* F_1_ hybrids with heterozygous *ORF5+* and *ORF5-*, which are functional alleles of aspartic protease (Chen et al., 2008; Yang et al., 2012), exhibit negative heterosis for seed setting rate. In contrast, hybrids carrying the *ORF5n*, which is assigned as a nonfunctional allele due to a 94 bp AV spanning the translation start site, alleviate the negative heterosis. Thus, the “nonfunctional” *ORF5n* allele shows significant function to heterosis for seed setting rate. Since inbred lines carrying any of the three alleles have normal embryo-sac fertility and seed setting rate, it is not appropriate to classify *ORF5n* as a dominant superior allele and *ORF5+*/*ORF5-* as recessive detrimental alleles under the dominance model. Similarly, heterozygous haplotypes of *OsBZR1* (haplotype_i/haplotype_JR2) outperformed homozygous ones for yield and multiple yield-related traits. Most importantly, our haplotype-based analysis of representative translocations and other SVs consistently indicates that the heterozygous state confers heterosis over the homozygous counterparts.

The overdominance model also posits the relationship between parental genetic disparity and the degree of heterosis. Previous studies using low- or high-density DNA markers have supported the close associations between genetic distance or genetic variants and heterosis of F_1_ hybrids in rice (Xiao et al., 1996; Zhang et al., 1996; Dan et al., 2014; Huang et al., 2015). We here offer a more comprehensive genomic variant picture based on high-quality genome assemblies and demonstrate that the number of genomic variants in parents are powerful predictors of heterosis for multiple agronomic traits. Crucially, the relationships differ between inter- and intra-subspecific hybrids, especially for heterosis of seed setting rate, tiller number per plant, and yield per plant. Based on the genome-wide heterozygosity of genomic variants, particularly SVs, and their correlations with heterosis, we support for the overdominance model and propose that heterozygosity of allelic/sequences underlies heterosis in rice.

Although we generated reference-level genome assemblies, telomere-to-telomere (T2T) genomes for all investigated inbred lines are needed to display the complete genomic landscape in subsequent heterosis studies. Concurrently, advanced variant detection and description tools should be developed to accurately characterize genomic variants in parental inbred lines. While large hybrid populations (often hundreds of F_1_ hybrids) are standard for robust heterosis studies, and hybridization and phenotyping are feasible in rice, generating hundreds of reference- or T2T-level genomes remains challenging for most research teams currently. Complete or half diallel-cross designs, which efficiently utilize parental genomic information, are practical strategies for deeper genomic investigation on rice heterosis. Additionally, the representativeness of parental inbred lines is important to outcomes. The 12 inbred lines used here capture only a portion of rice genetic diversity. The inclusion of accessions from *aus*, *tropical japonica*, *basmati*, and wild accessions (e.g. *rufipogon*) will provide updated genomic insights into rice heterosis.

The SVs detected in our study ranged from about 57 Mb to168 Mb, constituting approximately 14% to 42% of the genomes. These SVs, particularly subspecies-specific inversions and AVs: inversion_6 and AV_6, provide a structural basis for *indica*-*japonica* divergence and genetic variation among inbred lines. Importantly, specific SVs are associated with heterosis of key agronomic traits, highlighting the application potential of SVs in genomic prediction. As noted, the 94 bp AV in *S5-ORF5* and the complex PAVs linked to *OsBZR1* drive heterosis for seed setting rate and yield, respectively. We propose that SVs distinguishing *indica* from *japonica* or divergent haplotypes are candidates for precise engineering to harness strong inter-subspecific heterosis. The identified translocations, particularly intronic_4, splicing_6, and upstream_7 related to changes in gene expression levels, are important targets. Although precise modulation of genomic SVs remains challenging, we believe that advances in genome editing technologies, such as DualPE (Zhao et al., 2025), will elucidate function of SVs and facilitate the utilization of heterosis in crops. Prior to establishing precise editing methods for SVs, e.g., translocations, the effects of specific SVs on heterosis can be indirectly validated via a population genetic strategy. Firstly, two inbred lines carrying targeted SVs are crossed to generate heterozygous F_1_ hybrids and subsequent recombinant inbred lines. These recombinant inbred lines—typically requiring at least five generations of self-crossing to achieve phenotypic uniformity— are then crossed with a male-sterile line to produce F_1_ hybrids for heterosis quantification of corresponding traits. To classify genotypes/haplotypes, the recombinant inbred lines and the sterile line are analyzed using standard PCR assays targeting the SVs. Finally, the functional impact of single SVs can be assessed by comparing heterosis between heterozygous and homozygous genotypes/haplotypes.

Genomic SVs always induce gene expression changes (Alonge et al., 2020; Shang et al., 2022). Moving beyond individual loci, our transcriptomic analyses reveal that rice genomic SVs, especially large size inversions and AVs, not only correctly classify inbred lines by subspecies but also effectively explain the variance in heterosis through the cumulative, additive expression of multiple genes. These SVs are disproportionately important drivers of transcriptional changes, with translocations (e.g., translocations located in splicing, upstream, and intronic regions) outperforming others. We detect associations among SVs, expression changes, and heterosis based on seedling transcriptomes. However, many genes are expressed in specific tissues (e.g., young panicles), stages, or environments. In addition to gene expression, genomic SVs may be linked to DNA methylation levels, chromatin accessibility (Wang et al., 2024), and histone modifications (Qi et al., 2024). Therefore, the collection of diversified genomic-downstream data, including epigenomic, transcriptomic, and translational data, is essential to design optimal SV heterozygosity in generating rice heterosis.

## Materials and methods

### Plant materials

Twelve rice inbred lines (ILs) were selected from accessions reported in previous studies (Dan et al., 2014; Dan et al., 2016; Dan et al., 2020; Dan et al., 2024). Based on clustering results from metabolite profiles generated using GC- or LC-MS platforms (Dan et al., 2016; Dan et al., 2020), low-density SSR markers (Dan et al., 2014), and high-density SNPs or InDels (Dan et al., 2024), these 12 inbred lines are representatives of *indica* and *japonica* accessions. Plants of the selected inbred lines for genome sequencing were grown under long-day conditions, maintaining a 16-hour light/8-hour dark photoperiod at 30 °C in a plant culture room. The seedlings were sown in 96-well PCR plates (without bottoms) and placed in a plastic basin filled with Yoshida nutrient solution (pH = 5.5). After 20 days of growth, leaves from each genotype were harvested for experimentation. For full-length RNA sequencing of YB and Balilla, the plants were divided into four tissue parts: root, middle leaf (the youngest leaf), leaf without middle leaf, and stem.

Additionally, seedlings of the 12 rice inbred lines were placed in an intelligent breeding room for RNA sequencing. The environmental conditions were set to 28 °C with a 16-hour light/8-hour dark photoperiod and controlled humidity. Each inbred line occupied half of a PCR plate and was maintained in a single basin containing four liters of nutrient solution, which was changed every three days. Fifteen-day-old seedings (without root) were harvested and frozen in liquid nitrogen. The middle five plants from each row were pooled to create one biological replicate, with three biological replicates established for each genotype.

The 12 inbred lines were crossed using a complete diallel design. As previously detailed (Dan et al., 2024), a total of 17 agronomic traits were recorded for both the inbred lines and their corresponding F_1_ hybrids. These traits included grain yield per plant (YPP), grain number per panicle (GNP), seed setting rate (SSR), thousand-grain weight (TGW), plant height/tiller number (at seedling, elongation, flowering and maturation stages) (SSPH, ESPH, FSPH, MSPH and SSTN, ESTN, FSTN, MSTN), heading date (HD), tiller number per plant (TPP), panicle length (PL), and the number of primary and secondary branches per panicle (PBN and SBN). Better-parent heterosis was calculated using the equation: BPH = (F_1_ - P_high_)/P_high_, where P_high_ represents the value of the high parent. For the heading date, parents with shorter heading dates were designated as P_high_, and the calculated values were subsequently multiplied by -1.

### Genomic sequencing

SMRTbell libraries (∼17 kb) for each rice inbred line were constructed using the SMRTbell Express Template Prep Kit 3.0 and sequenced on the Pacbio Revio platform (Pacific Biosciences) at Frasergen Bioinformatics (Wuhan, China). The SMRTbell libraries were size-selected using the BluePippin system (Sage Science) and purified with a 1× concentration of SMRTbell cleanup beads. Library quality was assessed using Femto pulse and Qubit 3.0 (Life Technologies), resulting in an average of 12.31 Gb of HiFi reads (approximately 30.46× coverage of the genome) obtained for each inbred line.

Hi-C libraries for YB and Balilla were prepared and sequenced using the BGI PE150 platform, generating a total of 51.81 Gb and 49.27 Gb of reads (approximately 127.80× coverage of the genome) for the two samples, respectively. The raw data were filtered using Trimmomatic (v0.40) (Bolger et al., 2014) with parameters LEADING:3 TRAILING:3 SLIDINGWINDOW:4:15 MINLEN:15. The quality of the clean data was evaluated with default parameters using FastQC (Wingett and Andrews, 2018).

### RNA-sequencing

Full-length RNA sequencing technology was employed to aid in the annotation of genes in YB and Balilla. Total RNA was extracted from four tissues of the seedling samples, and equal amounts of RNA were pooled before constructing SMRTbell libraries. Full-length cDNA was synthesized using the SMARTer PCR cDNA Synthesis Kit (Clonetech) with the primer sequences “AAGCAGTGGTATCAACGCAGAGTACATGGGG” and “AAGCAGTGGTATCAACGCAGAGTAC”. The products were purified and size-filtered (>1 kb) prior to sequencing. The PacBio Sequel II platform at Frasergen Bioinformatics was used for sequence analysis, with raw data (an average of 36.14 Gb) filtered and preprocessed using SMRTlink (v10.1, --minLength=50). Iso-seq analysis was performed to obtain full-length and non-concatemer (FLNC) sequences. These FLNC sequences were mapped to reference genomes using GMAP (V19.06.10) (Wu and Watanabe, 2005), retaining those with the highest PID to determine gene structure.

To investigate the impact of structural variants on gene expression changes, RNA sequencing was also conducted to seedlings of the 12 rice inbred lines (with three biological replicates per genotype). Total RNA was extracted for each sample to construct a DNA library. The Illumina Hiseq PE150 platform at Frasergen Bioinformatics was selected to generate 150 bp paired-end reads, with more than 6 Gb of raw data per sample. The raw data were filtered using SOAPnuke (v2.1.0, - lowQual=20, -nRate=0.005, -qualRate=0.5) (Chen et al., 2018). Clean reads were aligned to YB genome using Hisat2 (v2.2.1, --no-unal) (Kim et al., 2019). Transcripts were assembled using StringTie (Pertea et al., 2015), and clean reads were mapped to reference transcripts using bowtie2 (v2.3.5, --sensitive -k 50 --no-mixed --no-discordant) (Langmead, 2010). The read count of each transcript was calculated using RSEM (v1.3.3) (Li and Dewey, 2011).

### Genome assembly

Draft genomes of the 12 rice inbred lines were assembled using HiFi reads with parameter -l3 via HiFiasm (v0.19.5) (Cheng et al., 2022). The clean Hi-C reads for YB and Balilla were aligned to contigs using Juicer (v1.6) (Durand et al., 2016). Contact matrices were generated with 3D-DNA (v180922, -r0) (Dudchenko et al., 2017), and corrections were applied using JuiceBox (v1.11.08). The genomes of the remained ten ILs were assembled based on the genomes of YB and Balilla using Ragtag (v2.1.0, scaffold) (Alonge et al., 2022). We assessed the accuracy, identity, and completeness of the 12 assemblies. The completeness of the assemblies was assessed using Benchmarking Universal Single-Copy Orthologue (v3.0.2, embryophyta_odb10) analysis (Simao et al., 2015). The long terminal repeats (LTR) assembly index (LAI) was calculated for each chromosome using LTR tools (Ou et al., 2018), including LTR_retriever (v2.9.0, -infinder ltrfinder.scn -inharvest ltrharvest.scn), LTR_FINDER (v1.0.7, -D 15000 -L 7000 -C -M 0.7), and GenomeTools (v1.6.2, ltrharvest -similar 70 -vic 10 -seed 20 -seqids yes -maxlenltr 7000 -maxtsd 6). HiFi reads were re-mapped to the assemblies with parameters “-ax map-hifi” using Minimap2 (v2.21) (Li, 2018). Additionally, consensus quality values and completeness of the assemblies were scored using a k-mer-based method with the Merqury pipeline (v1.3) (Rhie et al., 2020).

### Genome annotation

Repetitive sequences (including tandem repeats and transposable elements), protein-coding genes, and non-coding RNAs were annotated for the genome assemblies of YB and Balilla. Tandem repeats were identified using the Tandem Repeats Finder (TRF, v4.09.1, 2 7 7 80 10 50 2000 -d -h) (Benson, 1999). Transposable elements were annotated through an integrated approach combining do novo and homology-based (Repbase, www.girinst.org/repbase (Bao et al., 2015)) strategies. The de novo repeat library was constructed using LTR_FINDER (v1.0.7, -w 2 -C) (Xu and Wang, 2007) and RepeatModeler (v2.0.1) (Flynn et al., 2020). Transposable elements were detected using RepeatMasker (v4.1.2) (Tarailo-Graovac and Chen, 2009) with parameters “- nolow -no_is -norna -parallel 2”, searching against both the Repbase (Bao et al., 2015) and de novo repeat library. Additionally, RepeatProteinMask (1.36) (Tarailo-Graovac and Chen, 2009) with parameters “-engine ncbi -noLowSimple -pvalue 0.0001” was utilized to identify transposable elements in Repbase (Bao et al., 2015).

To annotate gene structure and function, we integrated three strategies: homology-based (using five genomes), de novo, and transcriptome-based (full-length RNA sequencing) predictions. In the homology-based approach, protein-coding sequences from the genomes of 9311 (Yu et al., 2002), Minghui63 (Zhang et al., 2016), Zhenshan97 (Zhang et al., 2016), R498 (Du et al., 2017), and Nipponbare (Shang et al., 2023) were compared to the YB and Balilla genomes using tblastn (v2.11.0+, -evalue 1e-05) (Gertz et al., 2006). Gene structures of the aligned sequences were predicted using Exonerate (v2.4.0, --model protein2genome) (Slater and Birney, 2005). De novo annotation was conducted using Augustus (v3.4.0, --noInFrameStop=true -- strand=both) (Stanke et al., 2006), GlimmerHMM (v3.0.4) (Majoros et al., 2004), and GeneMark (v4.65) (Borodovsky and McIninch, 1993) . The annotated gene sets were integrated to non-redundant sets using MAKER (v3.01.03) (Holt and Yandell, 2011), and further polished by incorporating full-length RNA sequencing results using PASA (v2.4.1) (Haas et al., 2003). RNA-sequencing results were aligned using HiSat2 (v.2.2.1) (Kim et al., 2019), and transcripts were assembled with StringTie (v.2.1.7) (Kovaka et al., 2019). Additionally, transcripts were de novo assembled using Trinity (v2.8.5) (Grabherr et al., 2011). Gene functional analysis was conducted with Diamond (v2.0.7, -c 1 -f 5 -e 1e-5 --very-sensitive -k 10) (Buchfink et al., 2021) against five databases: Non-Redundant (NR, ftp://ftp.ncbi.nlm.nih.gov/blast/db/FASTA/nr.gz), Kyoto Encyclopedia of Genes and Genomes (KEGG) (Kanehisa et al., 2014), Gene Ontology (GO) (Ashburner et al., 2000), TrEMBL (Boeckmann et al., 2003), and Swiss-Prot (Boeckmann et al., 2003). Protein domains were annotated using InterProScan (v5.50-84.0, --applications Pfam) (Jones et al., 2014) based on the InterPro database (Mitchell et al., 2015).

Non-coding RNAs, including rRNA, tRNA, snRNA, and miRNA, were also annotated for the two genomes. tRNA was identified using tRNAscan-SE (v2.0.9) (Lowe and Eddy, 1997), while rRNA was detected using RNAmmer (v1.2, -S euk - multi -m lsu,ssu,tsu) (Lagesen et al., 2007). miRNA and snRNA were annotated with Infernal (v1.1.4) (Nawrocki and Eddy, 2013) using Rfam (v14.6) (Griffiths-Jones et al., 2005). miRNA and rRNA were present in higher percentages within the genomes. The completeness of the genomic annotations was further assessed using BUSCO (v5.2.2, embryophyta_odb10) (Simao et al., 2015). Here, 98.5% and 98.6% of the gene sets in the two genomes contained complete BUSCOs in embryophyta.

For the annotation of the remaining ten genomes, Liftoff (Shumate and Salzberg, 2021) was employed. The YB genome was chosen as the reference genome for five *indica* accessions: R465, Mianhui725, 9311K, 610234, and Qianlijing. The Balilla genome served as the reference for five *japonica* accessions: R4115, C418, 110080, Wuyunjing8, and JR2.

### Pan-genome analysis

Protein coding genes of the 12 inbred lines were initially aligned using BLASTp (- evalue 1e-5) (Altschul et al., 1997). The BLAST results were processed with OrthoFinder (v2.3.12) (Emms and Kelly, 2019) to classify gene families. Gene families shared among all 12 inbred lines were defined as core gene families, while those identified in more than 90% of inbred lines were termed softcore gene families. Gene families specific to a single inbred line were classified as private gene families, and the remaining gene families were considered dispensable. Genes in the four types of gene families were further analyzed in the Pfam (Mistry et al., 2021), GO (Ashburner et al., 2000), and KEGG (Kanehisa et al., 2014) databases to determine their conservation.

### Identification of genomic variants

Genomic variants, including single nucleotide polymorphisms (SNPs), insertions and deletions (InDels) (2-50 bp), and structural variants (SVs), were detected in the 12 rice inbred lines, with the genome of YB selected as the reference genome and the remained 11 as query genomes. SNPs and InDels were detected using MUMmer4 (nucmer: -c 90 -l 40, delta-filter: -1, show-SNPs: -ClrTH) (Marcais et al., 2018). SVs, including presence-absence variants (PAVs), inversions, translocations, and copy number variants (CNVs), were detected with the assistance of SyRI (v1.4, nucmer: -c 90 -l 40 -t 16 ref.fa query.fa, delta-filter: -1 -i 90 -l 100 ref_query.delta, show-coords: -THrd) (Goel et al., 2019). The distribution of SNPs, InDels, and SVs was analyzed across various genomic regions or functions, including: 1. Upstream (1 kb regions before the gene start codon); 2. Downstream (1 kb regions after the gene stop codon); 3. Intergenic regions; 4. UTR3 (3’ untranslated regions); 5. UTR5 (5’ untranslated regions); 6. Splicing junction regions (< 10 bp); 7. Intronic regions; 8. Exonic regions; 9. Synonymous or nonsynonymous mutants; 10. Frameshift or nonframeshift insertions/deletions; 11. Stopgain (gain of stop codons); 12. Stoploss (loss of stop codons). Variants simultaneously located in two regions, such as UTR5 and UTR3, were showed as Region 1;Region 2. The functions of SNPs, InDels, and SVs was annotated with ANNOVAR (Wang et al., 2010).

### Pan-SV analysis

A pan-SV analysis was conducted for SVs of the 11 query genomes and the results were further used for constructing graph-based genome using Vg (v1.43) (Garrison et al., 2018). For SVs excluding PVs, an SV was assigned when the ratio of overlapping to total locations in two inbred lines exceeded 50%. For PVs, a PV was designated when the distance between PV locations in two inbred lines was less than 5 bp and the length ratio between the shorter and longer samples was greater than 50%. Circles illustrating genomic variants on chromosomes were generated using Circos (Krzywinski et al., 2009). Genetic relatedness among the 12 inbred lines was visualized using Cytoscape (Reimand et al., 2019).

### Construction of phylogenetic trees

A phylogenetic tree for 223 rice inbred lines was constructed using genotypic data based on 33 pairs of single sequence repeat markers from a previous study (Dan et al., 2024). Phylogenetic trees based on the number of differential SNPs, InDels, and CNVs among the 12 rice inbred lines were built using the neighbour-joining method with MEGA (v10.2.6) (Stecher et al., 2020). The number of SNPs, InDels, and CNVs of the inbred lines were retrieved from a previous report (Dan et al., 2024).

For phylogenetic analysis of *Phr1* and the *S5* loci, gene sequences from reference inbred lines were obtained from the GenBank database. The *Phr1* sequences of Nipponbare and 9311 corresponded to accession number DQ532376 and DQ532391, respectively (Yu et al., 2008). For *S5-ORF3-5* (Chen et al., 2008; Yang et al., 2012), the Balilla sequences were retrieved under accession number JX138498, JX138502, EU889294. Gene sequences from the 12 newly assembled inbred lines were extracted and aligned using Jalview (version 2.11.4.1) with default clustal parameter s (Waterhouse et al., 2009). Phylogenetic trees were constructed with the neighbour-joining method. Genotypes of inbred lines were assigned based on previous reports (Chen et al., 2008; Yu et al., 2008; Yang et al., 2012). Gene structures were visualized using the exon-intron graphic maker (http://wormweb.org/exonintron).

To investigate genomic variants of *OsBZR1*, especially SVs in promoter region, sequences (including 6,345 bp before the ATG) of the 12 ILs were used for analysis. In sequence alignment, the mafft combined with E-INS method was chosen. The dendrogram tree of the 12 ILs based on phenotypic data of seven traits were implemented using MetaboAnalyst 6.0 (Pang et al., 2022), with Spearman and ward as the distance measure and clustering algorithm, respectively.

### Phenol reaction experiment

Seeds from 14 inbred lines were harvested at maturity and stored at -20 °C. For each inbred line, three seeds were chosen and treated with a 2% (volume/volume) phenol solution at 37 °C. Changes in hull colour of the seeds were recorded after 1, 2, 3, 5, 7, 31, 62, 93, and 124 days of treatment. After 7 days of treatment, the seed were placed at room temperature.

### Normalization and differential analysis of transcriptomics data

Read counts of 53,564 genes in each experimental sample were used for data normalization and differential analyses through MetaboAnalyst 5.0 (Pang et al., 2022). The interquantile range was selected to filter genes, with a threshold set at 40%. Sum, none, and auto-scaling were chosen for sample normalization, data transformation, and data scaling, respectively. Differential analysis among the 12 inbred lines was implemented using the ANOVA module in MetaboAnalyst 5.0 (Pang et al., 2022), with a raw *P*-value cutoff set at 0.001 for post-hoc analysis. Principal component analysis (PCA) and heatmap results for the 12 inbred lines were obtained through corresponding modules on the website. PERMANOVA was conducted in PCA and the permutation number was 999. Euclidean for distance measure and ward for clustering method were set in heatmaps. Differential genes were selected from the original read count table and averaged across the three biological replicates. The averaged data were further filtered and normalized for each genotype with the above parameters. Due to the limited number of genes in inversions and absence variants, no filtering was applied to these genes. The genes in three large AVs, including AV_6, AV_8, and AV_11, were analyzed. Despite AV_8 and AV_11 being larger in size, the genes in these AVs were rarely expressed in the seedling samples. Only genes in AV_6 was further analyzed.

To investigate the relationship between expression level of *OsBZR1* (*Os07g0580500*) and heterosis, gene expression data and seedling length (15-day old seedlings) of six pairs of reciprocal hybrids and corresponding parental inbred lines were obtained from a previous study (Dan et al., 2024). The transcriptomic data were Log2-normalized using NetworkAnalyst 3.0 (Zhou et al., 2019). Since several genotypes had six biological replicates, seeding length of the first three replicates was used for the calculation of better-parent heterosis and Spearman correlation analysis. To determine inheritance patterns of *OsBZR1*, the reciprocal hybrids from two inbred lines were treated as single genotype. Average expression levels of *OsBZR1* were calculated between each pair of inbred lines and were treated as additive expression levels. Gene expression levels were compared for the parents, F_1_ hybrids, and additive expression levels. Inheritance patterns were assigned based on previous rules (Dan et al., 2024).

### Partial least square (PLS) analysis

PLS analysis was conducted to investigate the relationship between inheritance patterns of expressed genes and better-parent heterosis of 17 agronomic traits using SPSS 20.0. The maximum number of latent factors was set to 50. The different inheritance patterns were manually calculated based on normalized gene expression levels using equations outlines in a previously study (Dan et al., 2024). The genetic effects included: 1. Additive effect: average levels of the female (P_f_) and male (P_m_) parents; 2. Partially dominant effects (PD): partially dominant _0.3 (PD_0.3) = P_m_+0.3×(P_f_-P_m_) and partially dominant _0.7 (PD_0.7) = P_m_+0.7×(P_f_-P_m_); 3. Dominant effects: expression levels of the female (dominant_female) and male (dominant_ male) parents. The model with the highest R^2^ (adjusted) was selected as the final model for each trait.

PLS was also used to study the relationship between differentially expressed genes in 12 inbred lines and the number of parental genomic variants. Additive effect of the differential genes was used as predictive variables to predict the number of unique variants between each two inbred lines.

### Statistical analysis

Boxplots were created using BoxPlotR (Spitzer et al., 2014). Analysis of variance (ANOVA) with the least significant difference for post hoc comparisons was performed using SPSS 20.0. This analysis compared the following parameters: lengths of genes, lengths of coding sequences, lengths of exons, lengths of introns, GC content of genes, GC content of coding sequences, number of exons, and number of introns across seven genomes (including two newly assembled genomes, YB and Balilla, along with five representative rice genomes). Additionally, we assessed the number of genomic variants and the degrees of better-parent heterosis for 17 traits in F_1_ hybrids from both intra- and inter-subspecies. The adjusted R^2^ values of partial least square models for heterosis across traits were also compared. An independent samples *t*-test (weighted) was conducted to evaluate the number of correlations between unique and common genomic variants and heterosis. SPSS was also employed to conduct Spearman correlation to investigate the relationships between different types of unique genomic variants and between genomic variants and heterosis.

## Acknowledgements

We thank C. Zhu for funding of our rice research. We thank J. Bao, L. Rong, and S. Zheng from the Frasergen Bioinformatics for their help in genomics and transcriptomics analyses.

## Author contributions

Z.D. conceived and designed the project. Z.D. and Y.C. prepared rice seedling samples, conducted genomic, transcriptomic, phenotypic, and statistical analyses, and interpretated the results. W.H. collected the rice accessions. Z.D. and W.H. constructed the diallel cross population and recorded the phenotypes. Z.D., Y.C., and W.H. wrote the manuscript.

## Supplementary data

The following materials are available in the online version of this article.

**Supplementary Figure S1.** Neighbour-joining tree of 223 rice accessions.

**Supplementary Figure S2.** Heterosis of 16 agronomic traits in F_1_ hybrids from intra- or inter-subspecies.

**Supplementary Figure S3.** Contact matrices and collinearity diagrams for genomes of YB and Balilla.

**Supplementary Figure S4.** Long terminal repeat assembly index (LAI) for 12 chromosomes of 12 rice accessions.

**Supplementary Figure S5.** Comparison of two newly assembled genomes with five established rice genomes.

**Supplementary Figure S6.** Genomic comparisons between Mianhui725, 610234, and YB.

**Supplementary Figure S7.** Genomic comparisons between 9311K, R465, and YB.

**Supplementary Figure S8.** Genomic comparisons between Qianlijing, C418, and YB.

**Supplementary Figure S9.** Genomic comparisons between R4115, Wuyunjing8, and YB.

**Supplementary Figure S10.** Genomic comparisons between 110080, JR2, and YB.

**Supplementary Figure S11.** Genomic variants displayed on the genome of YB.

**Supplementary Figure S12.** Genetic relatedness of 12 rice inbred lines based on different types of genomic variants.

**Supplementary Figure S13.** Number of genomic variants for F_1_ hybrids from intra- and inter-subspecies.

**Supplementary Figure S14.** Correlations between the number of structural variants and heterosis of intra- and inter-subspecific F_1_ hybrids.

**Supplementary Figure S15.** Dendrogram of 12 inbred lines based on phenotypic data of seven agronomic traits.

**Supplementary Figure S16.** Inheritance patterns of *OsBZR1* across F_1_ hybrids.

**Supplementary Figure S17.** Transcriptomic additive and partially dominant effects contribute to heterosis of 17 traits.

**Supplementary Figure S18.** Associations between expression levels of genes in inversion_6 and heterosis.

**Supplementary Figure S19.** Associations between expression levels of genes in inversion_8 and heterosis.

**Supplementary Figure S20.** Genomic structure of two AVs and the prediction of heterosis with AV_6.

**Supplementary Figure S21.** Adjusted R^2^ for different types of SNPs.

**Supplementary Figure S22.** Adjusted R^2^ for different types of InDels.

**Supplementary Figure S23.** Genomic structure and expression patterns of five translocations.

**Supplementary Figure S24.** Associations between genes in translocation_4/translocation_12.2 and heterosis.

**Supplementary Figure S25.** Adjusted R^2^ for different types of AVs.

**Supplementary Figure S26.** Adjusted R^2^ for different types of CNVs.

**Supplementary Figure S27.** Adjusted R^2^ for different types of PVs.

**Supplementary Figure S28.** Adjusted R^2^ for different types of inversions.

**Supplementary Figure S29.** Heterozygosity of genomic variants in F_1_ hybrids.

**Supplementary Table S1.** Genome assembly profile of 12 rice inbred lines.

**Supplementary Table S2.** Consensus quality values and completeness of the 12 assemblies.

**Supplementary Table S3.** Hi-C profile of two rice genomes.

**Supplementary Table S4.** Annotation of two rice genomes.

**Supplementary Table S5.** Genomic variants of each pair of inbred lines and heterosis of their F_1_ hybrids.

## Data availability

Raw reads of PacBio SMRT sequencing and RNA-sequencing of the 12 rice inbred lines have been deposited in the NCBI database under BioProject accession PRJNA1178436. Genome assemblies of the 12 rice inbred lines are available from Figshare repository with DOI: 10.6084/m9.figshare.28397960. Source data are available with this paper.

