## Supplementary figures for "Structural variations contribute to subspeciation and yield heterosis in rice"

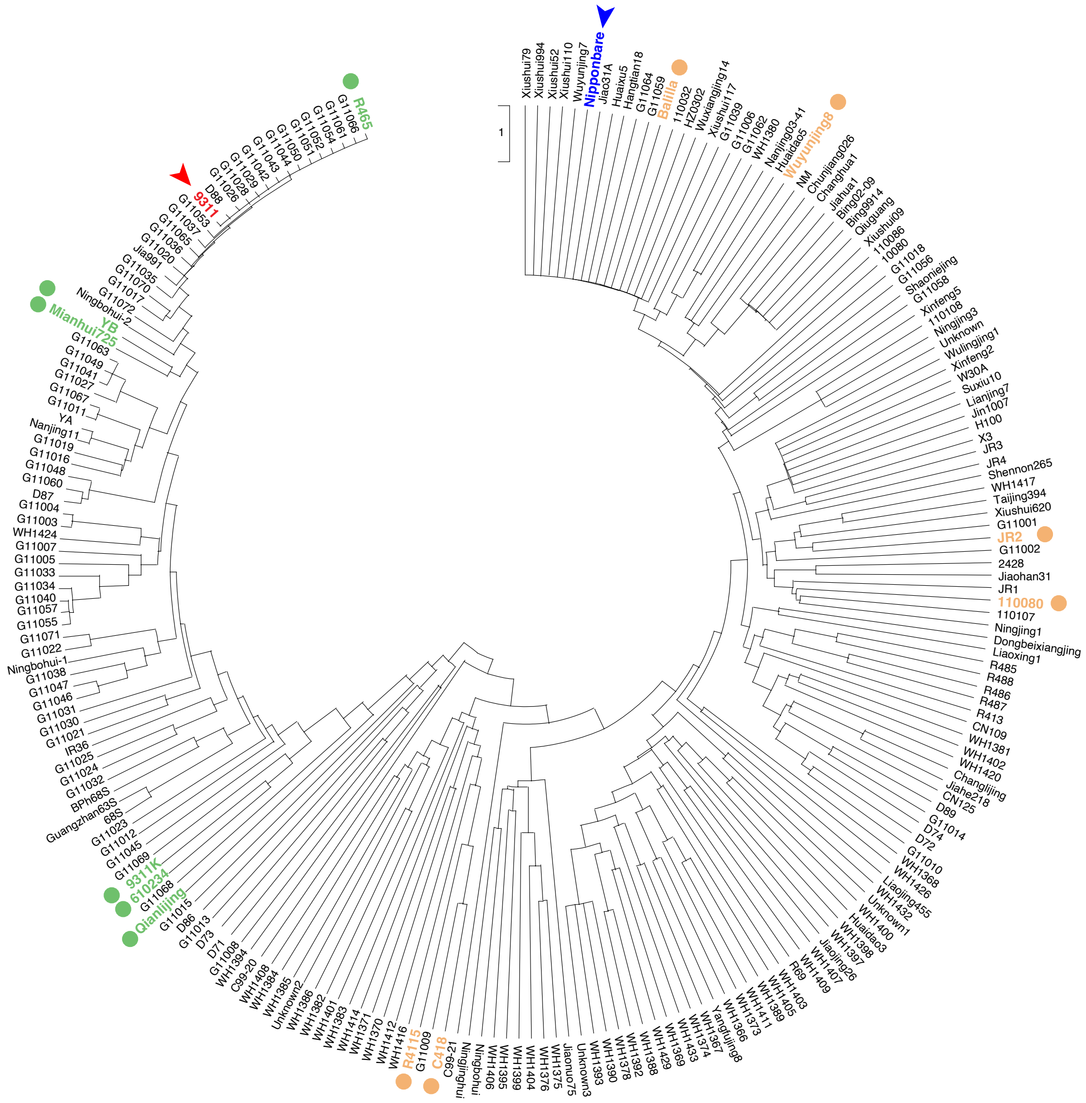


**Supplementary Figure S1.** Neighbour-joining tree of 223 rice accessions. A total of 33 pairs of single sequence repeat DNA markers were used to genotype these inbred lines. The 12 accessions utilized in this study are indicated in green for *indica* and brown for *japonica*, respectively. Two representative accessions, 9311 (*indica*) and Nipponbare (*japonica*), are marked in red and blue, respectively.

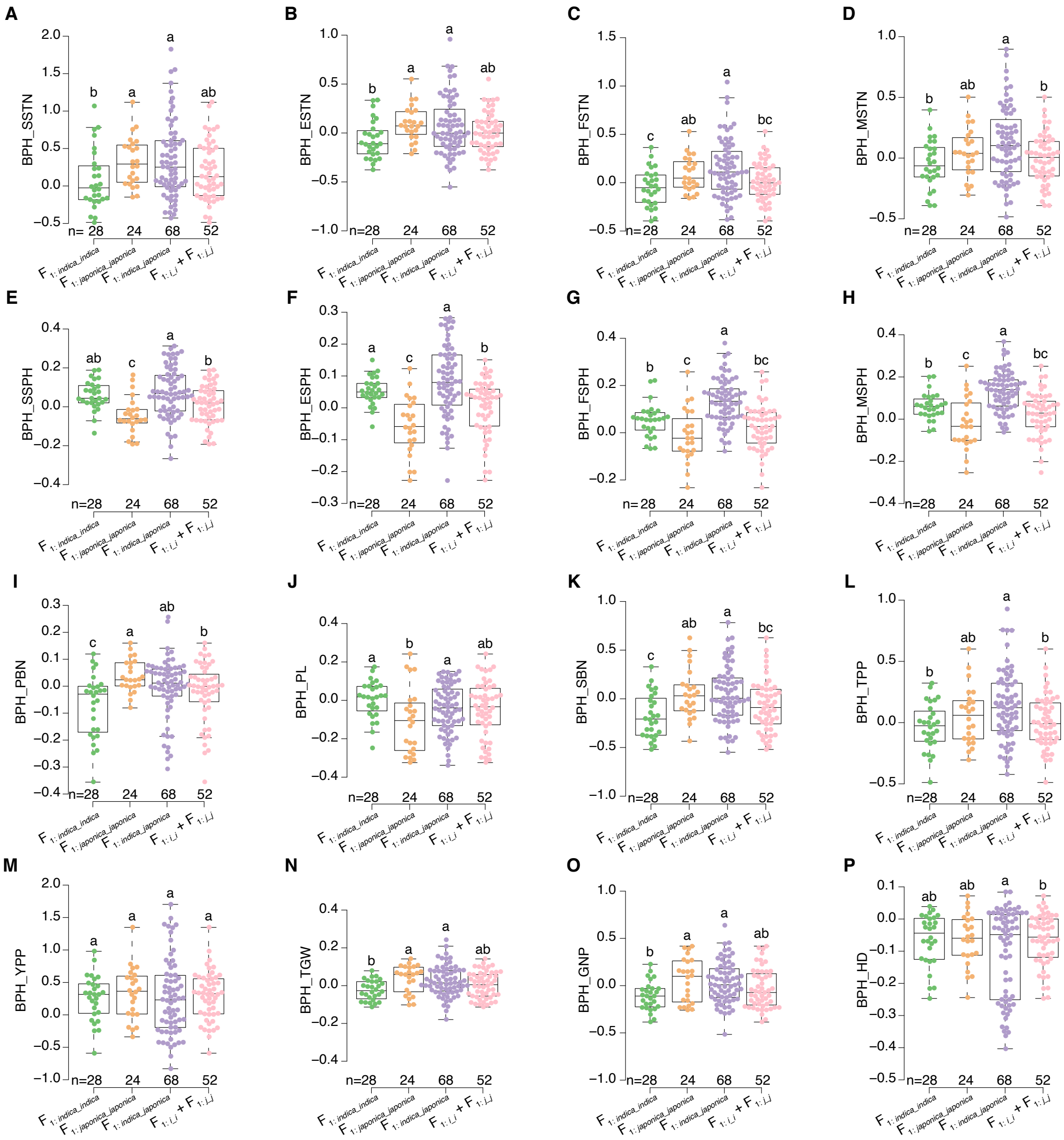

**Supplementary Figure S2.** Heterosis of 16 agronomic traits in $F_1$ hybrids from intra- or inter-subspecies. Better-parent heterosis (BPH) of 16 traits was compared for *indica-indica* (*i_i*), *japonica-japonica* (*j_j*), *indica-japonica*, and the pooled intra-subspecific $F_1$ hybrids. The traits include: seedling stage tiller number (SSTN; **A**), elongation stage tiller number (ESTN; **B**), flowering stage tiller number (FSTN; **C**), maturation stage tiller number (MSTN; **D**), seedling stage plant height (SSPH; **E**), elongation stage plant height (ESPH; **F**), flowering stage plant height (FSPH; **G**), maturation stage plant height (MSPH; **H**), primary branch number (PBN; **I**), panicle length (PL; **J**), secondary branch number (SBN; **K**), tiller number per plant (TPP; **L**), yield per plant (YPP; **M**), thousand-grain weight (TGW; **N**), grain number per panicle (GNP; **O**), and heading date (HD; **P**). Letters indicate results of ANOVA with LSD in post-hoc test (*P*-value < 0.05).

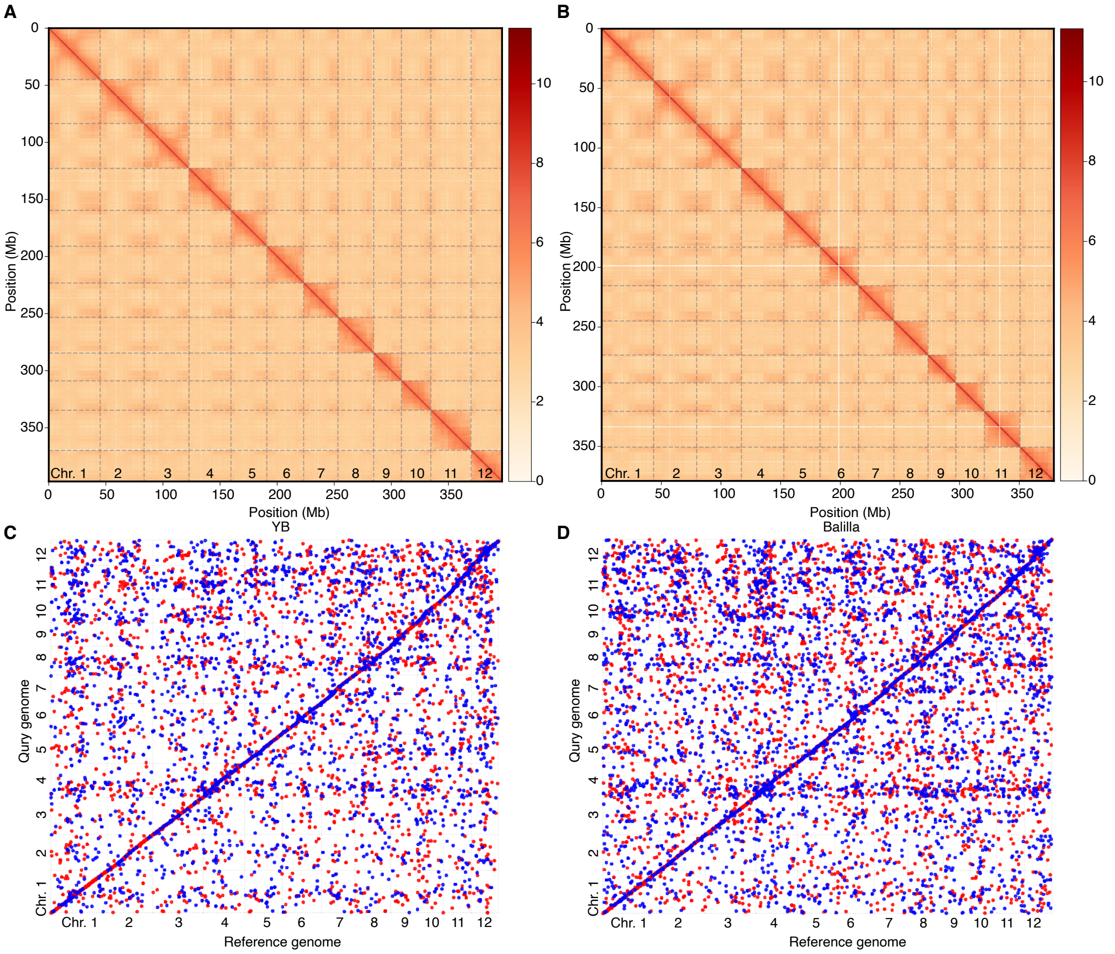


**Supplementary Figure S3.** Contact matrices and collinearity diagrams for genomes of YB and Balilla. Hi-C data were utilized to aid genome assembly for YB and Balilla. Contact matrices display interactions for YB (**A**) and Balilla (**B**). Collinearity analysis illustrates matching results for YB (**C**) and Balilla (**D**).

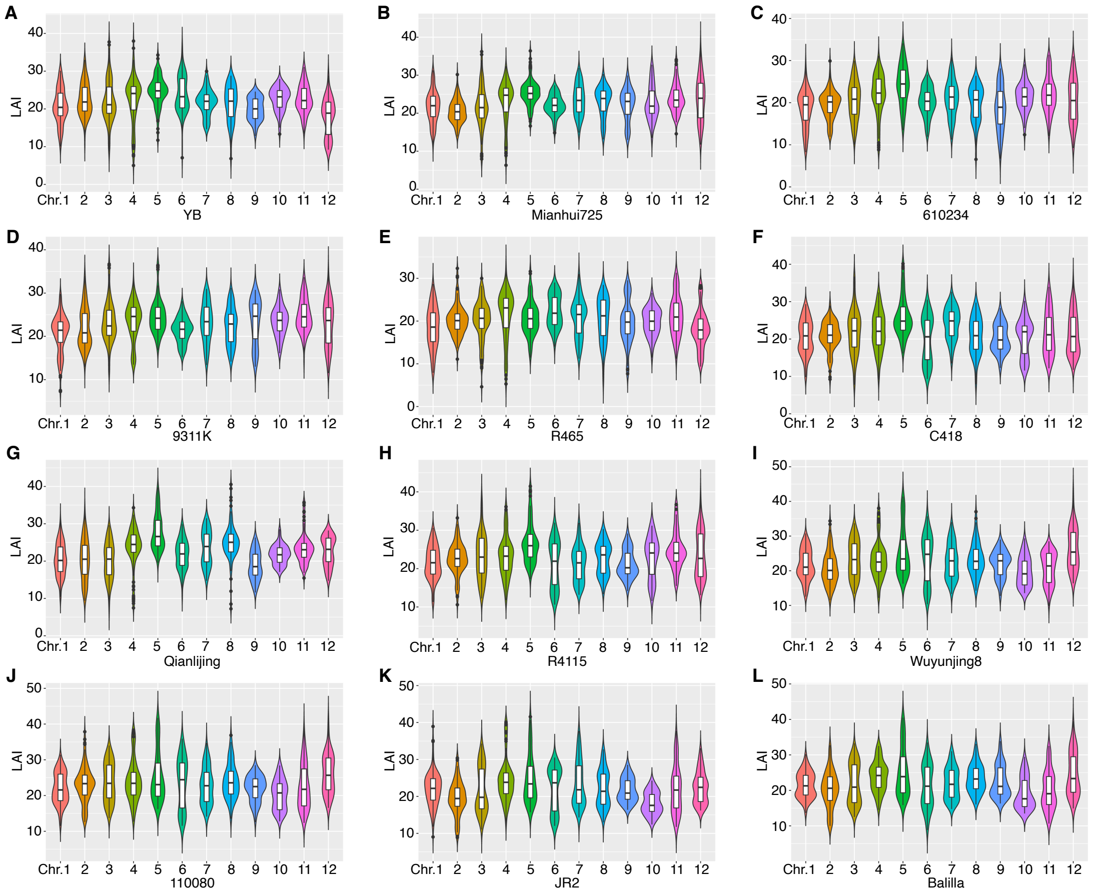

**Supplementary Figure S4.** Long terminal repeat assembly index (LAI) for 12 chromosomes of 12 rice accessions. LAI was calculated for each chromosome of 12 inbred lines. Results are shown for YB (**A**), Mianhui725 (**B**), 610234 (**C**), 9311K (**D**), R465 (**E**), C418 (**F**), Qianlijing (**G**), R4115 (**H**), Wuyunjing8 (**I**), 110080 (**J**), JR2 (**K**), and Balilla (**L**).

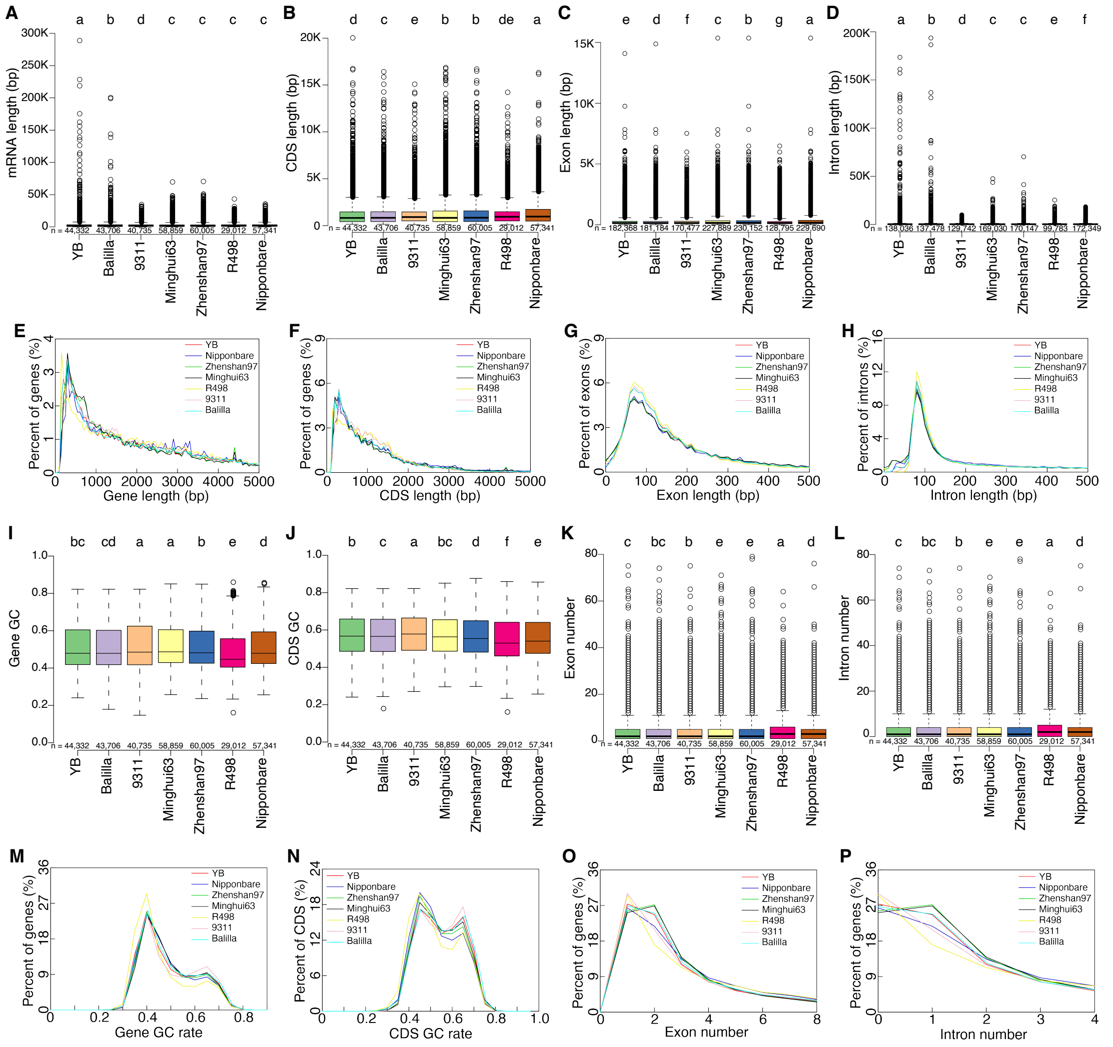


**Supplementary Figure S5.** Comparison of two newly assembled genomes with five established rice genomes. **A)** Length of mRNA. **B)** Length of coding sequences (CDS). **C)** Length of exons. **D)** Length of introns. **E)** Percentages of genes with varying lengths. **F)** Percentages of genes with different lengths of coding sequences. **G)** Percentages of exons with varying lengths. **H)** Percentages of introns with varying lengths. **I)** GC content of genes. **J)** GC content of coding sequences. **K)** Number of exons per gene. **L)** Number of introns per gene. **M)** Percentages of genes with different GC content. **N)** Percentages of coding sequences with different GC content. **O)** Percentages of genes with varying number of exons. **P)** Percentages of genes with varying number of introns. The newly assembled genomes are for YB (*indica*) and Balilla (*japonica*). The five established genomes include 9311 (*indica*), Minghui63 (*indica*), Zhenshan97 (*indica*), R498 (*indica*), and Nipponbare (*japonica*). Letters indicate results of ANOVA with LSD in post-hoc test ($P$-value < 0.05).

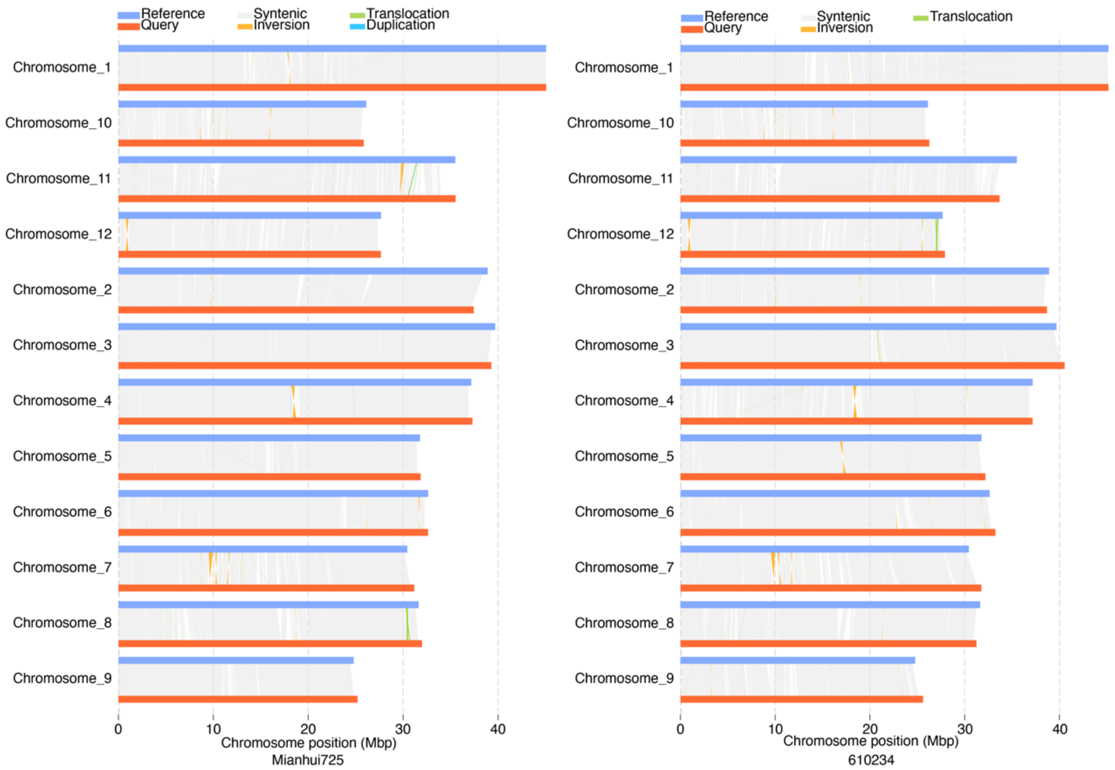


**Supplementary Figure S6.** Genomic comparisons between Mianhui725, 610234, and YB. The genome of YB serves as the reference genome, while Mianhui725 and 610234 are the query genomes. Syntenic regions are connected with grey lines, and inversions, translocations, and duplications between the genomes are shown.

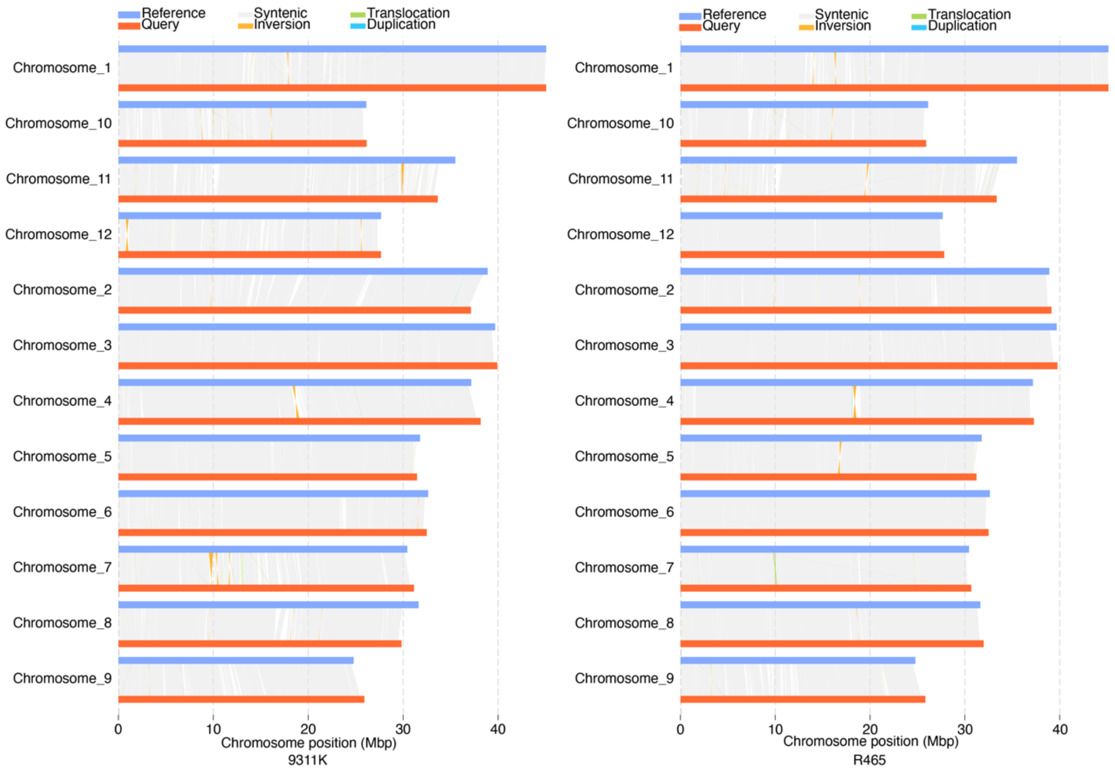


**Supplementary Figure S7.** Genomic comparisons between 9311K, R465, and YB. YB is the reference genome, with 9311K and R465 as the query genomes. Syntenic regions are connected with grey lines, and inversions, translocations, and duplications between the genomes are shown.

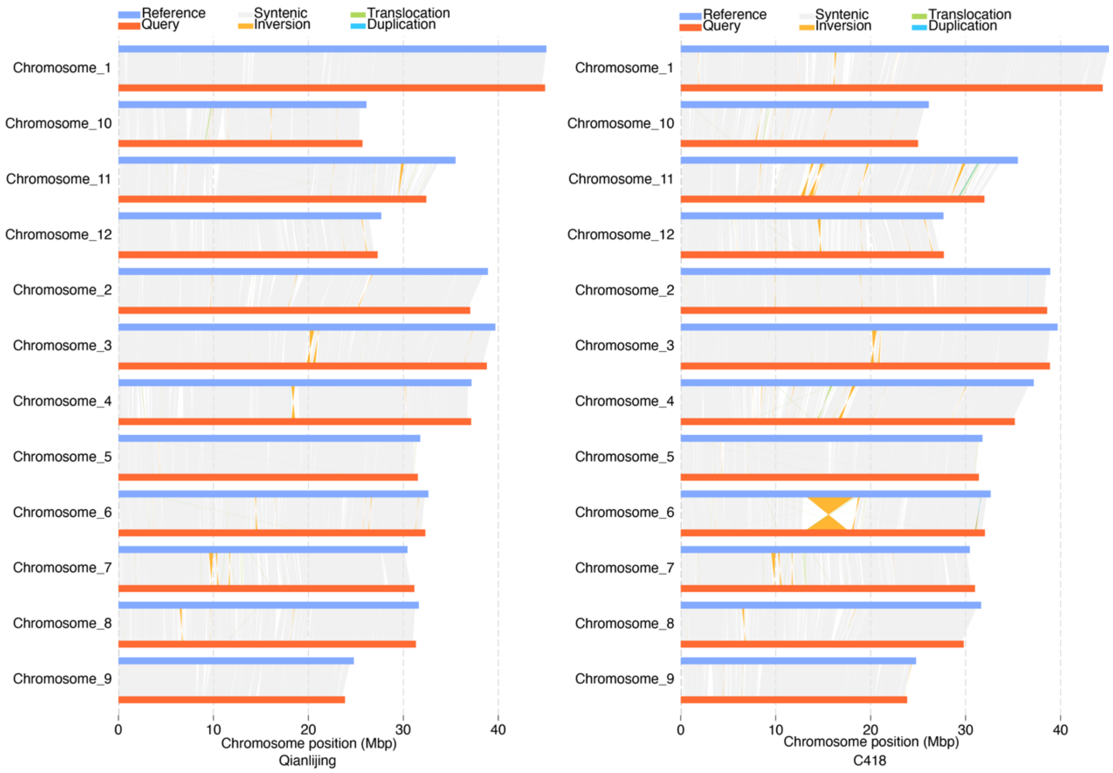


**Supplementary Figure S8.** Genomic comparisons between Qianlijing, C418, and YB. YB serves as the reference genome, with Qianlijing and C418 as the query genomes. Syntenic regions are connected with grey lines, and inversions, translocations, and duplications between the genomes are shown.

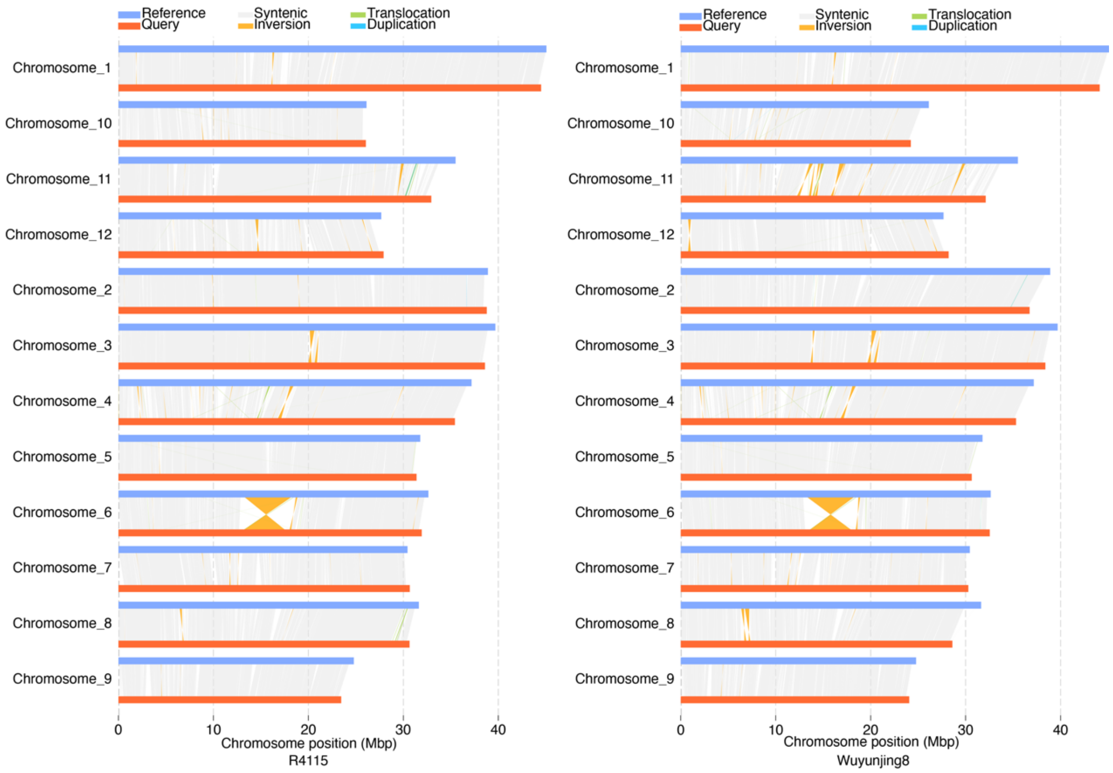


**Supplementary Figure S9.** Genomic comparisons between R4115, Wuyunjing8, and YB. YB is the reference genome, while R4115 and Wuyunjing8 are the query genomes. Syntenic regions are connected with grey lines, and inversions, translocations, and duplications between the genomes are shown.

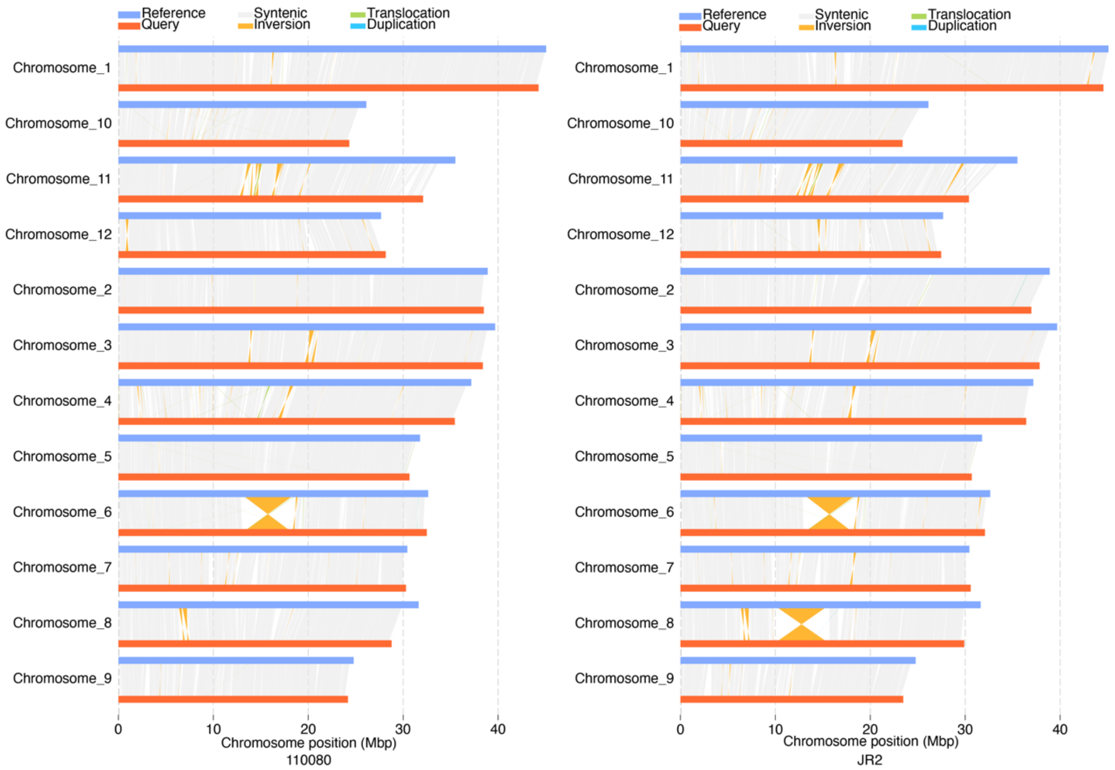


**Supplementary Figure S10.** Genomic comparisons between 110080, JR2, and YB. The genome of YB serves as the reference genome, with 110080 and JR2 as query genomes. Syntenic regions are connected with grey lines, and inversions, translocations, and duplications between the genomes are shown.

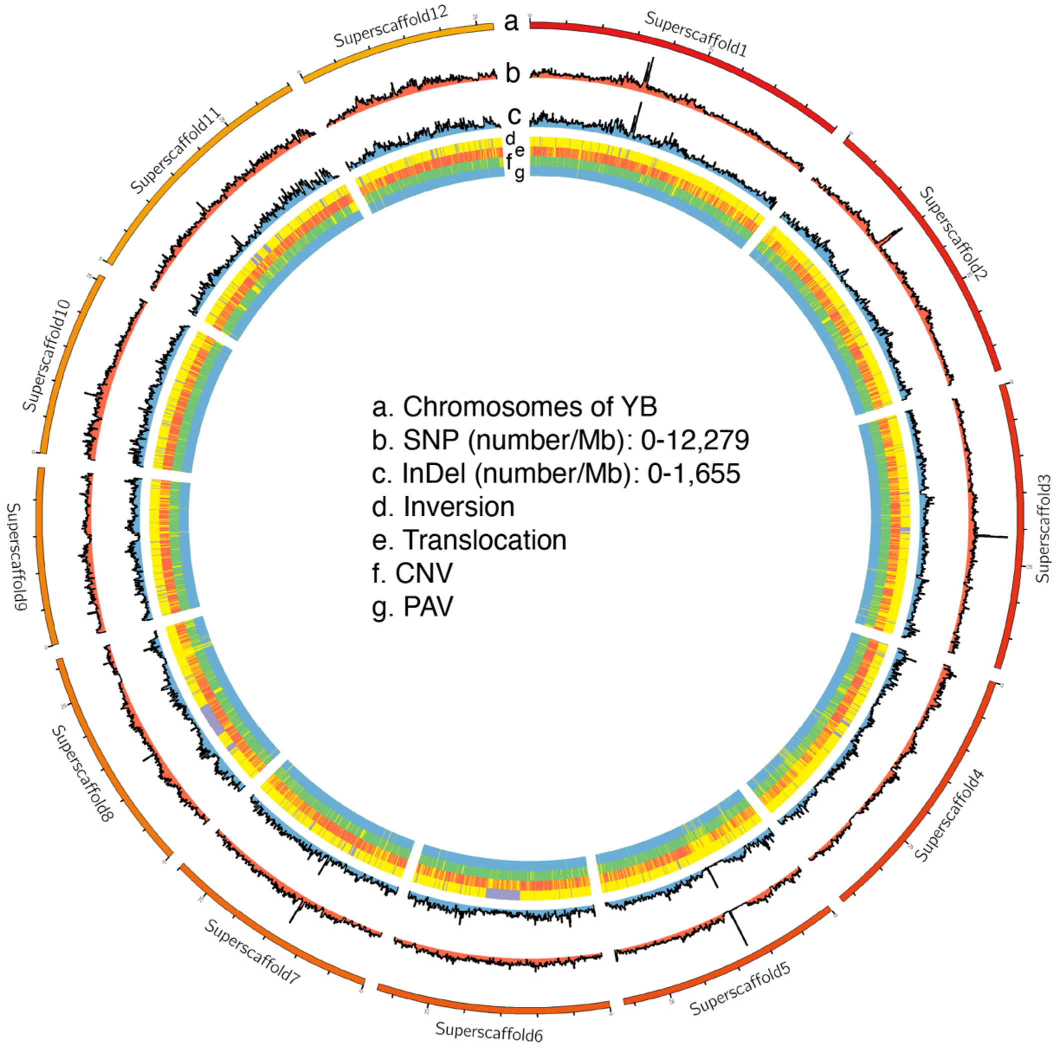


**Supplementary Figure S11.** Genomic variants displayed on the genome of YB. The genomic variants, including single nucleotide polymorphisms (SNPs), insertions and deletions (InDels), inversions, translocations, copy number variants (CNVs), and presence/absence variants (PAVs), are displayed across the 12 chromosomes of YB. Superscaffold represents chromosome.

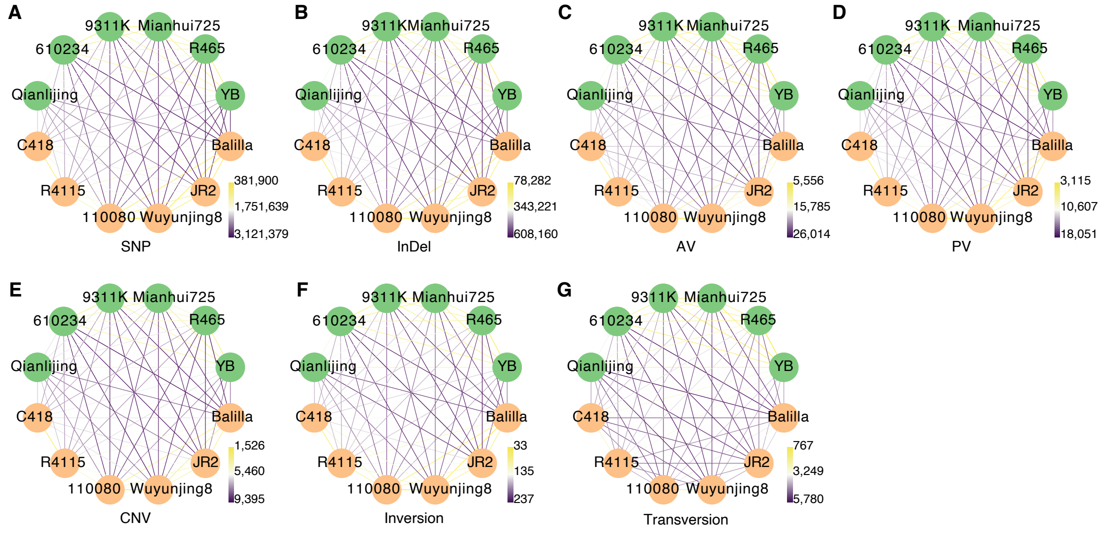


**Supplementary Figure S12.** Genetic relatedness of 12 rice inbred lines based on different types of genomic variants. The number of unique genomic variants for each pair of inbred lines is used to illustrate genetic relatedness. Genomic variants include single nucleotide polymorphisms (SNPs; **A**), insertions and deletions (InDels; **B**), absence variants (AVs; **C**), presence variants (PVs; **D**), copy number variants (CNVs; **E**), inversions (**F**), and translocations (**G**).

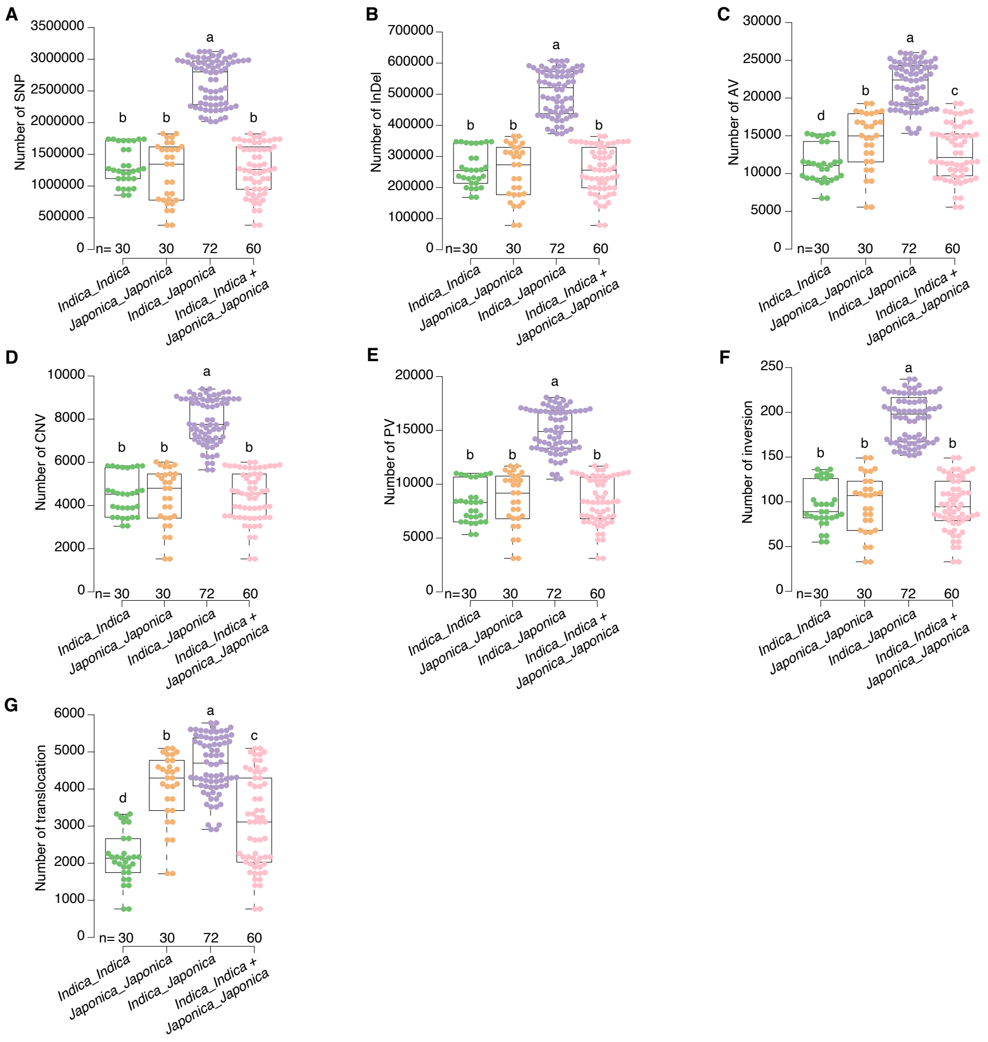


**Supplementary Figure S13.** Number of genomic variants for $F_1$ hybrids from intra- and inter-subspecies. Genomic variants include: single nucleotide polymorphisms (SNPs; **A**), insertions and deletions (InDels; **B**), absence variants (AVs; **C**), copy number variants (CNVs; **D**), presence variants (PVs; **E**), inversions (**F**), and translocations. ANOVA with LSD in post-hoc test (*P*-value < 0.05) was performed for comparison.

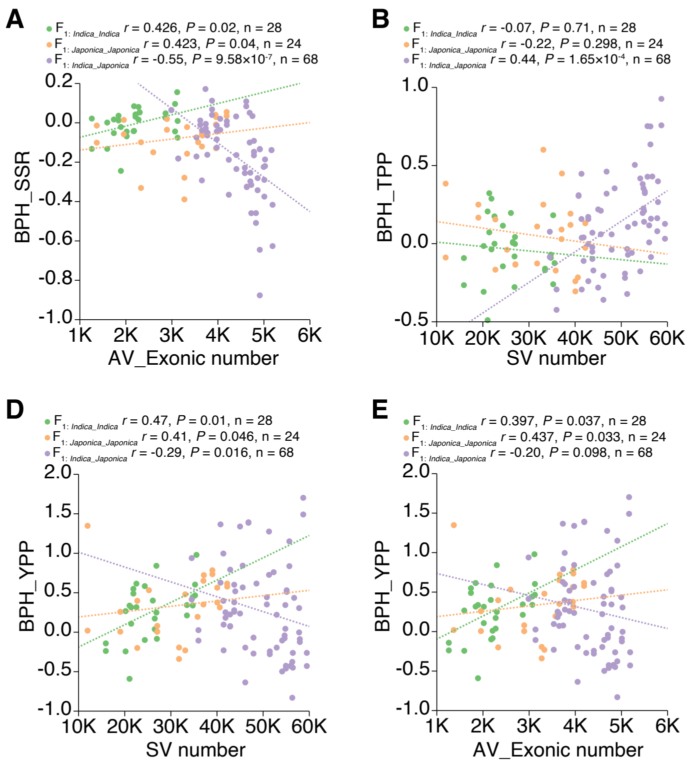


**Supplementary Figure S14.** Correlations between the number of structural variants and heterosis of intra- and inter-subspecific $F_1$ hybrids. Better-parent heterosis for seed setting rate (BPH_SSR; **A**), tiller number per plant (BPH_TPP; **B-C**), and yield per plant (BPH_YPP; **D-E**) were investigated. The number of structural variants (SVs) and absence variants (AVs) in exonic regions was analyzed, with *P*-values indicating Spearman correlation.

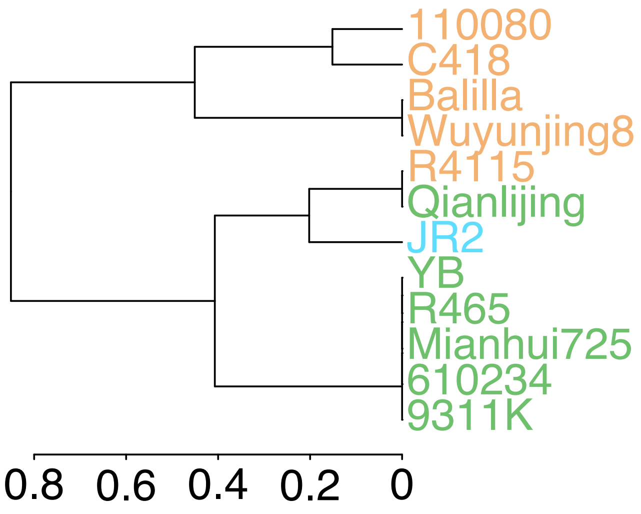


**Supplementary Figure S15.** Dendrogram of 12 inbred lines based on phenotypic data of seven agronomic traits. These traits include yield per plant, tiller number per plant, grain number per panicle, thousand-grain weight, panicle length, maturation stage plant height, and seed setting rate.

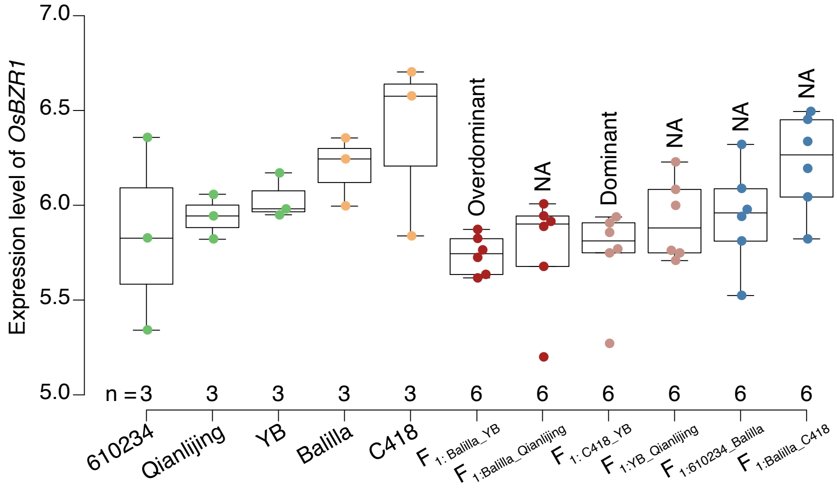


**Supplementary Figure S16.** Inheritance patterns of *OsBZR1* across $F_1$ hybrids. Expression levels of *OsBZR1* were compared for parents, $F_1$ hybrids, and additive effect. NA represents unclassifiable patterns.

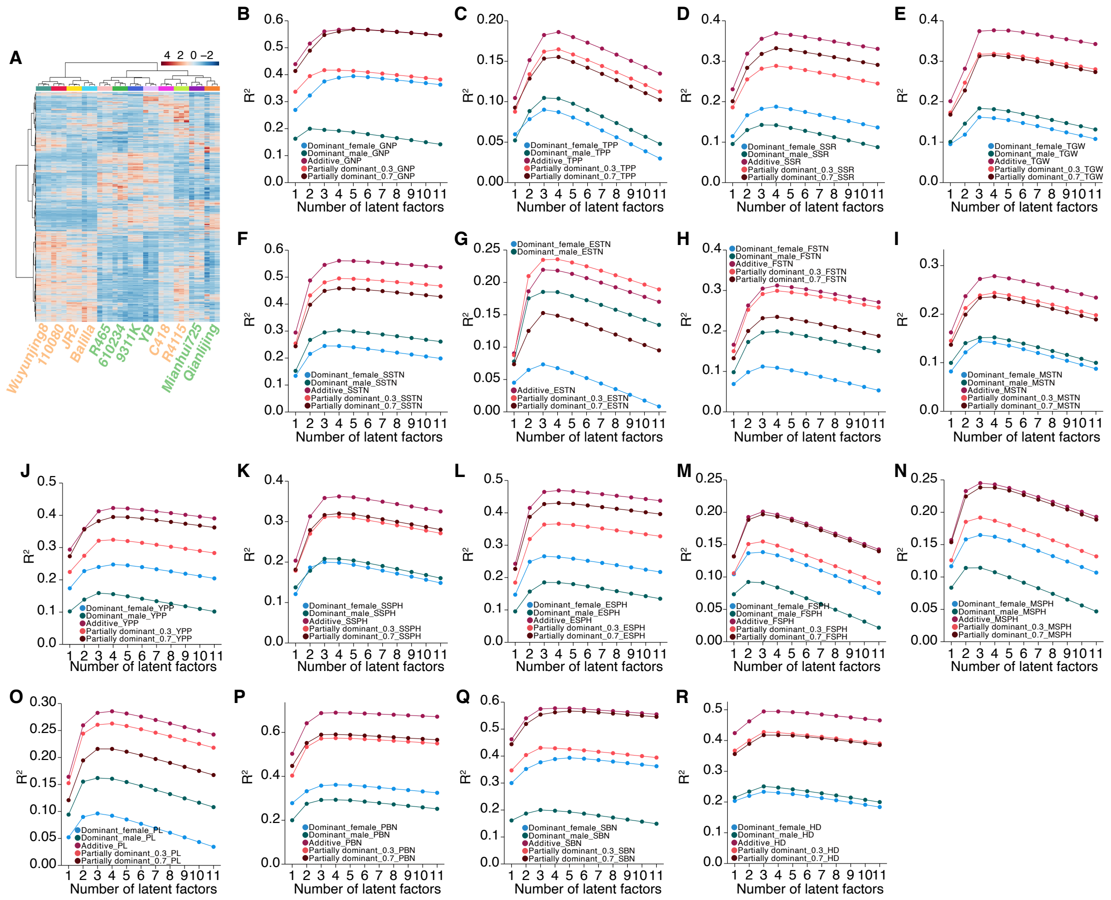

**Supplementary Figure S17.** Transcriptomic additive and partially dominant effects contribute to heterosis of 17 traits. **A)** Heatmap of 12 rice inbred lines based on their differentially expressed genes (DEGs). **B-R)** Changes in adjusted $R^2$ values with varying number of latent factors. Partial least square-based predictive models were built for better-parent heterosis (BPH) of 17 traits. The traits include grain number per panicle (GNP; **B**), tiller number per plant (TPP; **C**), seed setting rate (SSR; **D**), thousand-grain weight (TGW; **E**), seedling stage tiller number (SSTN; **F**), elongation stage tiller number (ESTN; **G**), flowering stage tiller number (FSTN; **H**), maturation stage tiller number (MSTN; **I**), yield per plant (YPP; **J**), seedling stage plant height (SSPH; **K**), elongation stage plant height (ESPH; **L**), flowering stage plant height (FSPH; **M**), maturation stage plant height (MSPH; **N**), panicle length (PL; **O**), primary branch number (PBN; **P**), secondary branch number (SBN; **Q**), and heading date (HD; **R**). Different inheritance patterns of DEGs, including dominant_female, dominant_male, additive, partially dominant_0.3, and partially dominant_0.7, served as the predictive variables.

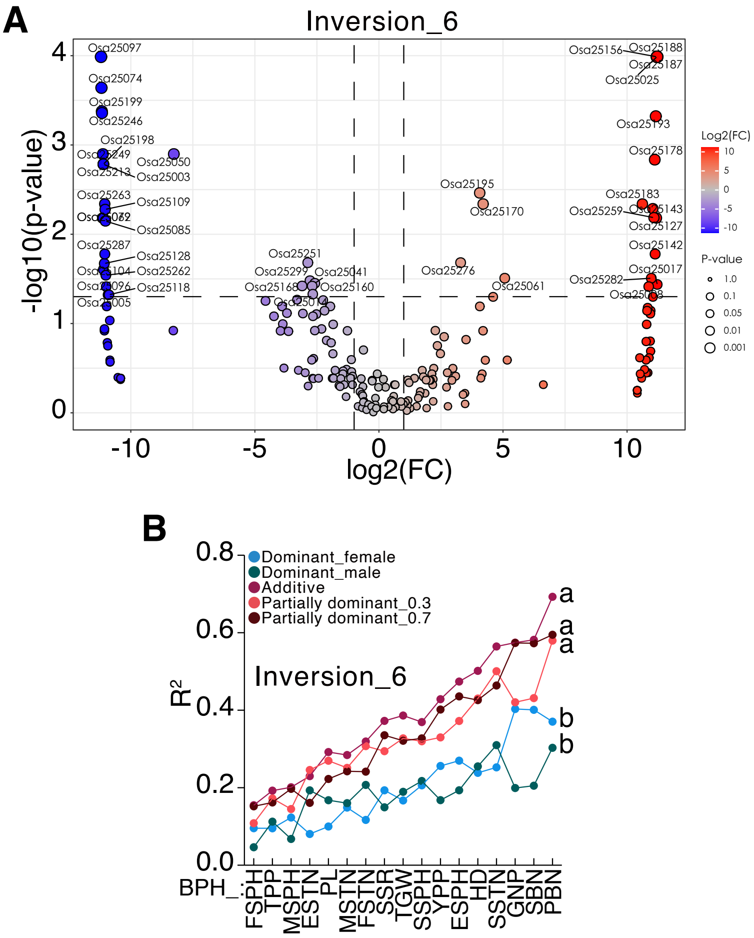


**Supplementary Figure S18.** Associations between expression levels of genes in inversion_6 and heterosis. **A)** Volcano plot of differentially expressed genes between *indica* and *japonica* for inversion_6. **B)** Adjusted $R^2$ values of partial least square models for better-parent heterosis of 17 traits. Different inheritance patterns of genes from inversion_6 served as predictive variables. The model with the highest $R^2$ value was selected as the final model for heterosis of each trait. Letters denote comparison results derived from ANOVA with LSD in post-hoc test (*P*-value < 0.05).

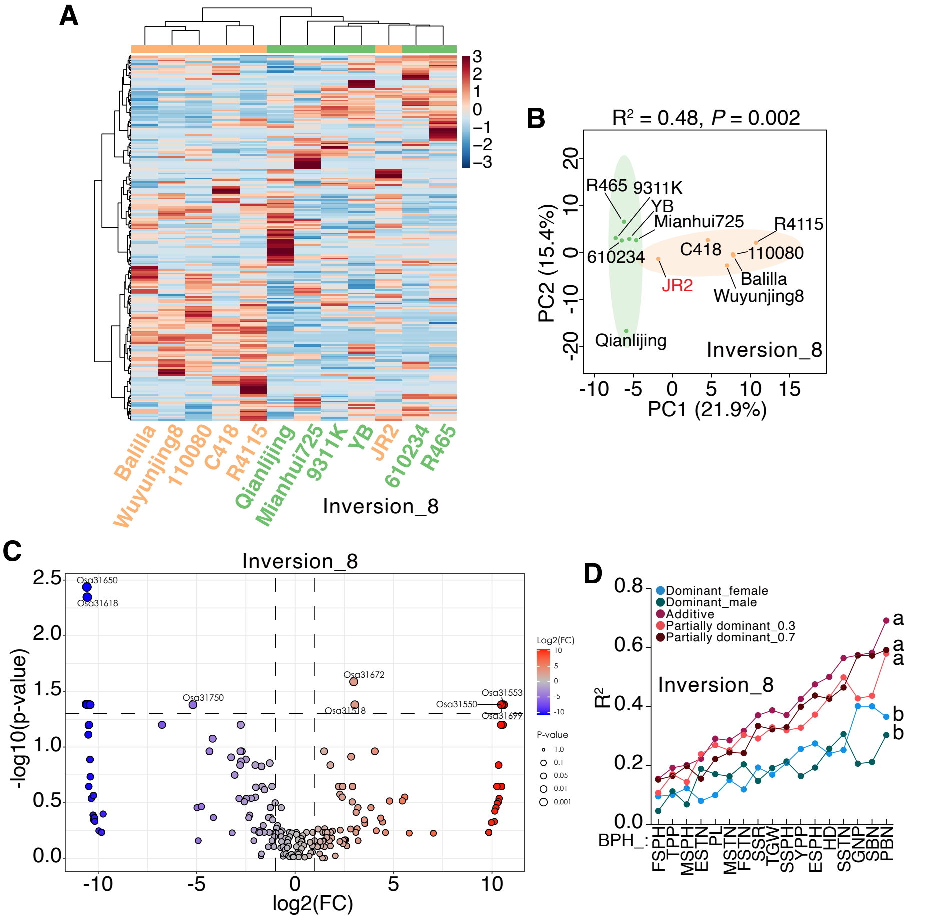


**Supplementary Figure S19.** Associations between expression levels of genes in inversion_8 and heterosis. **A)** Heatmap for expression levels of genes in inversion_8 across 12 inbred lines. **B)** PCA score plot of 12 inbred lines based on gene expression levels of inversion_8. *P*-value indicates the result from PERMANOVA. **C)** Volcano plot of differential genes between *indica* and *japonica* for inversion_8. **D)** Adjusted $R^2$ values for predicting heterosis based on expression levels of genes in inversion_8. Letters denote comparison results derived from ANOVA with LSD in post-hoc test ($P$-value < 0.05).

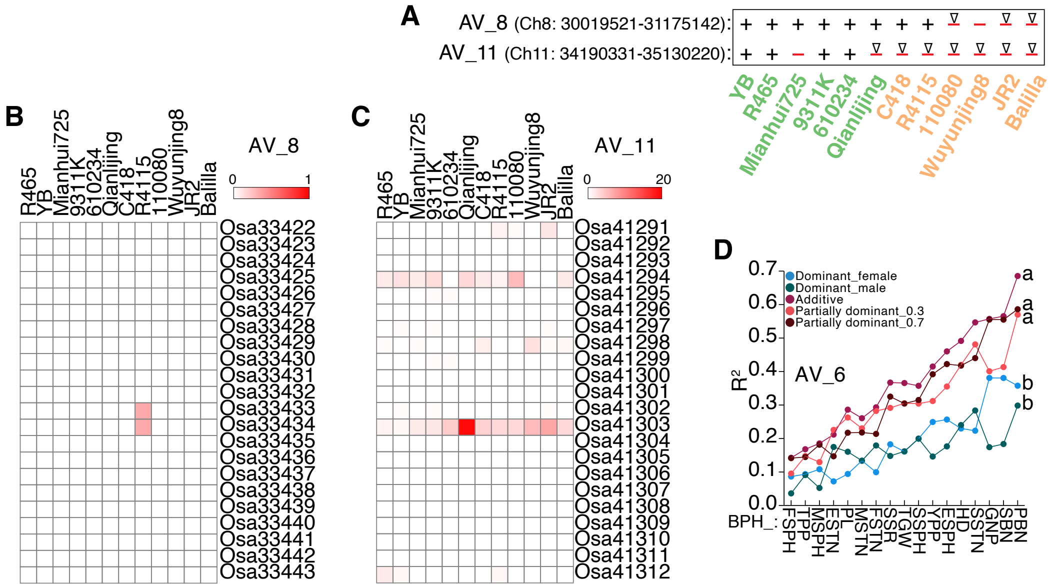


**Supplementary Figure S20.** Genomic structure of two AVs and the prediction of heterosis with AV_6**. A)** Genomic structure of AV_8 and AV_11 in 12 inbred lines. **B-C)** Expression patterns of genes in AV_8 and AV_11. **D)** Adjusted $R^2$ values for predicting heterosis based on expression levels of genes in AV_6. Letters denote comparison results derived from ANOVA with LSD in post-hoc test (*P*-value < 0.05).

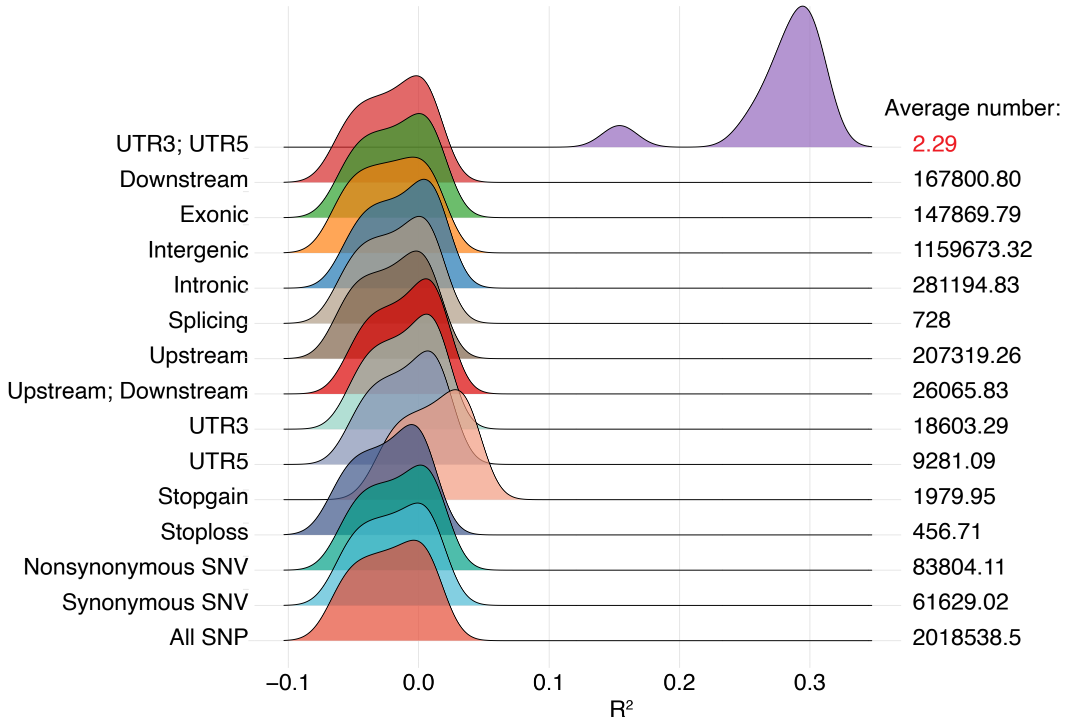


**Supplementary Figure S21.** Adjusted $R^2$ for different types of SNPs. The number of parental unique SNPs were predicted with additive effect of differentially expressed genes in 12 inbred lines using partial least square model. The SNPs were classified into different types based on their location or function.

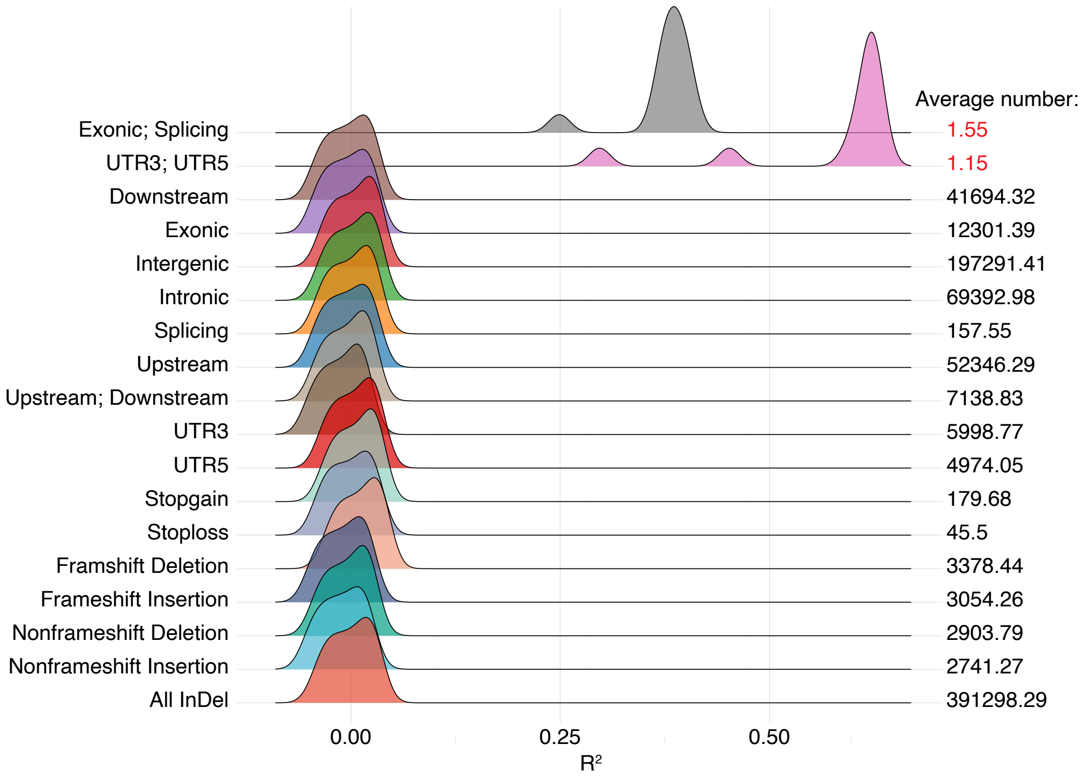


**Supplementary Figure S22.** Adjusted $R^2$ for different types of InDels. The number of parental unique InDels were predicted with additive effect of differentially expressed genes in 12 inbred lines using partial least square model. The InDels were classified into different types based on their location or function.

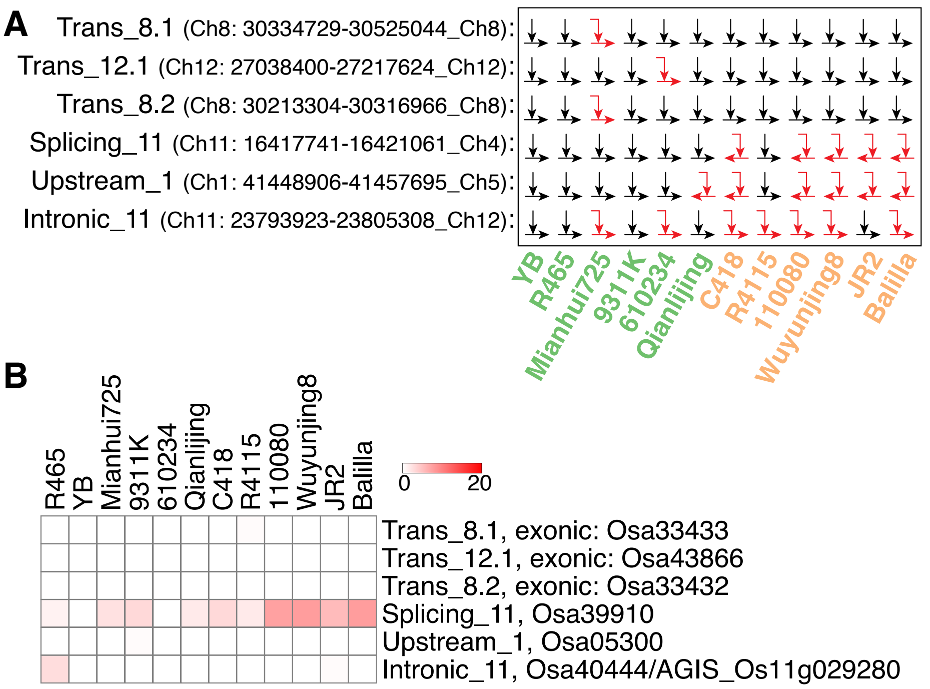


**Supplementary Figure S23.** Genomic structure and expression patterns of five translocations. **A)** Genomic structure of five translocations in 12 inbred lines. **B)** Expression patterns of five translocations in 12 inbred lines. Trans represents translocation.

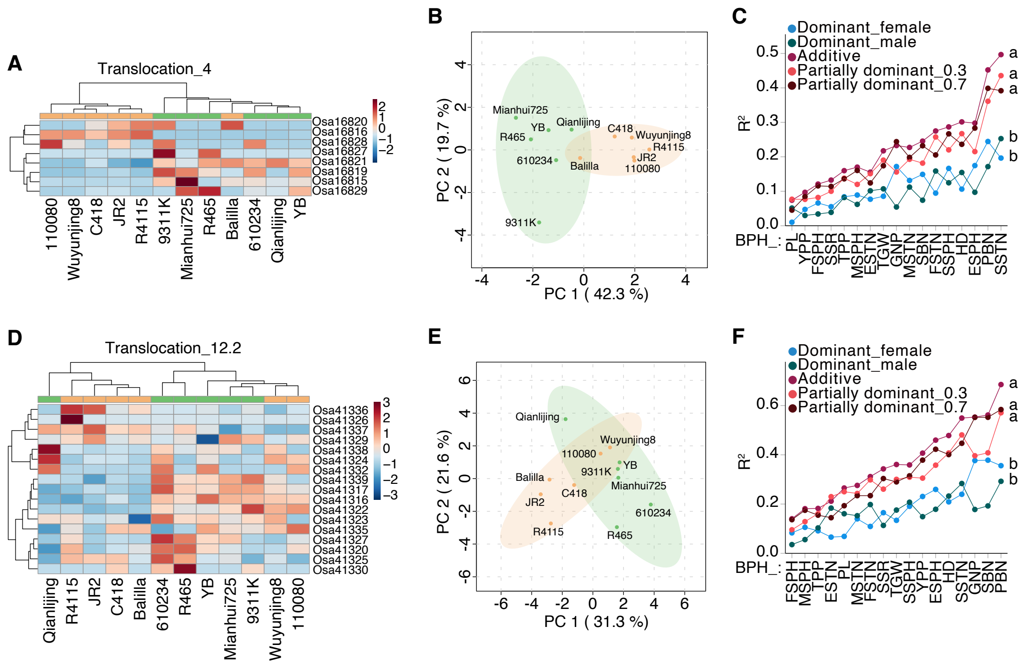


**Supplementary Figure S24.** Associations between genes in translocation_4/translocation_12.2 and heterosis. **A, D)** Heatmaps for expression levels of genes in translocation_4 and translocation_12.2 across 12 inbred lines. **B, E)** PCA score plot of 12 inbred lines based on expressed genes in translocation_4 and translocation_12.2. **C, F)** Adjusted $R^2$ values for predicting heterosis based on expression levels of genes in translocation_4 and translocation_12.2. Letters denote comparison results derived from ANOVA with LSD in post-hoc test ($P$-value < 0.05).

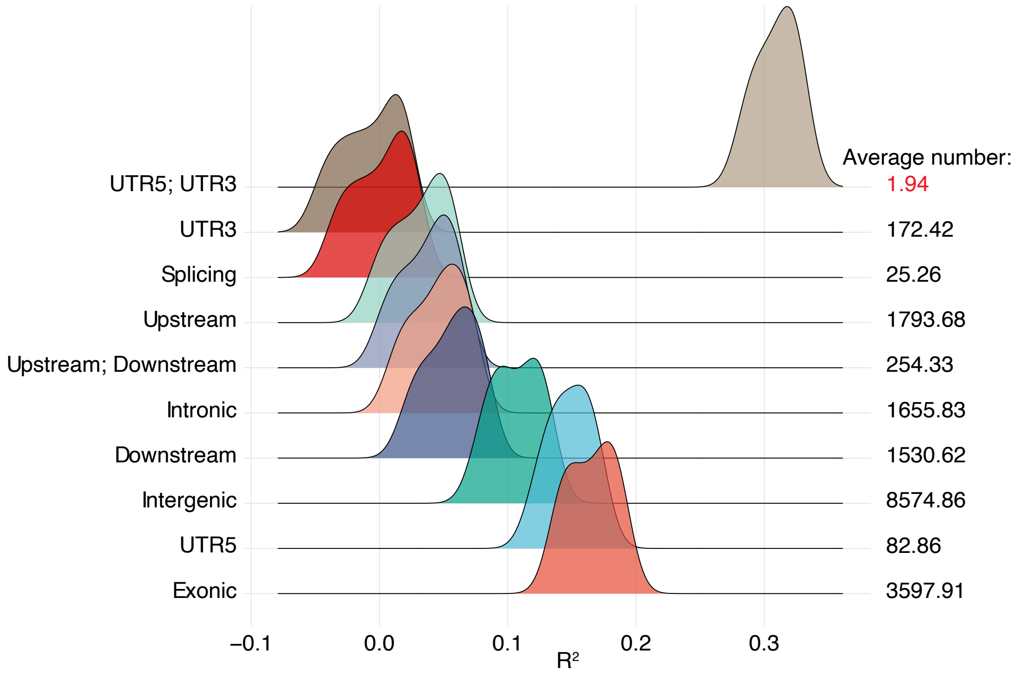


**Supplementary Figure S25.** Adjusted $R^2$ for different types of AVs. The number of parental unique AVs were predicted with additive effect of differentially expressed genes in 12 inbred lines using partial least square model. The AVs were classified into different types based on their location.

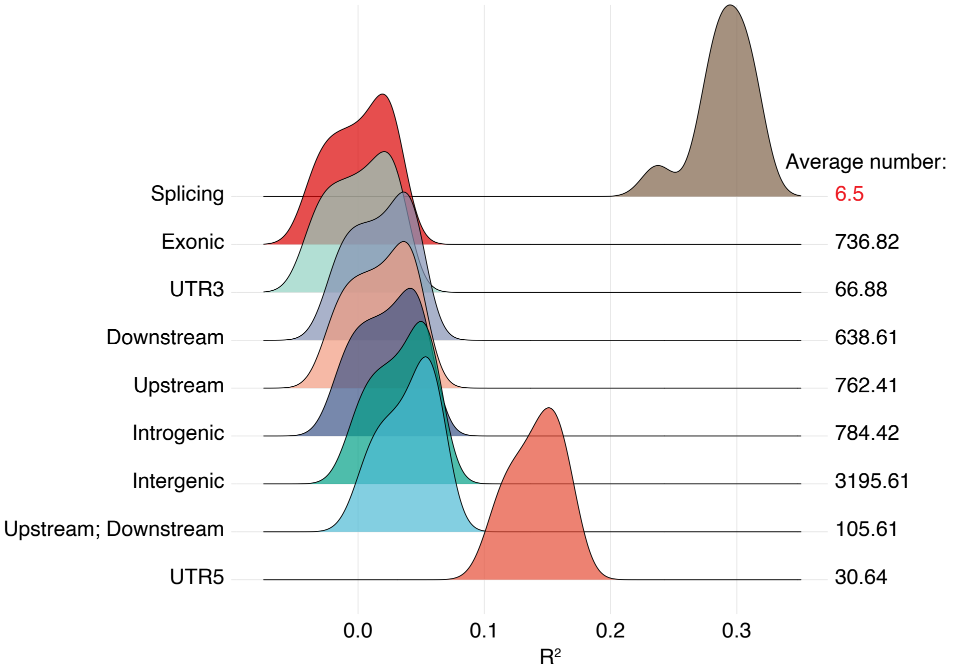


**Supplementary Figure S26.** Adjusted $R^2$ for different types of CNVs. The number of parental unique CNVs were predicted with additive effect of differentially expressed genes in 12 inbred lines using partial least square model. The CNVs were classified into different types based on their location.

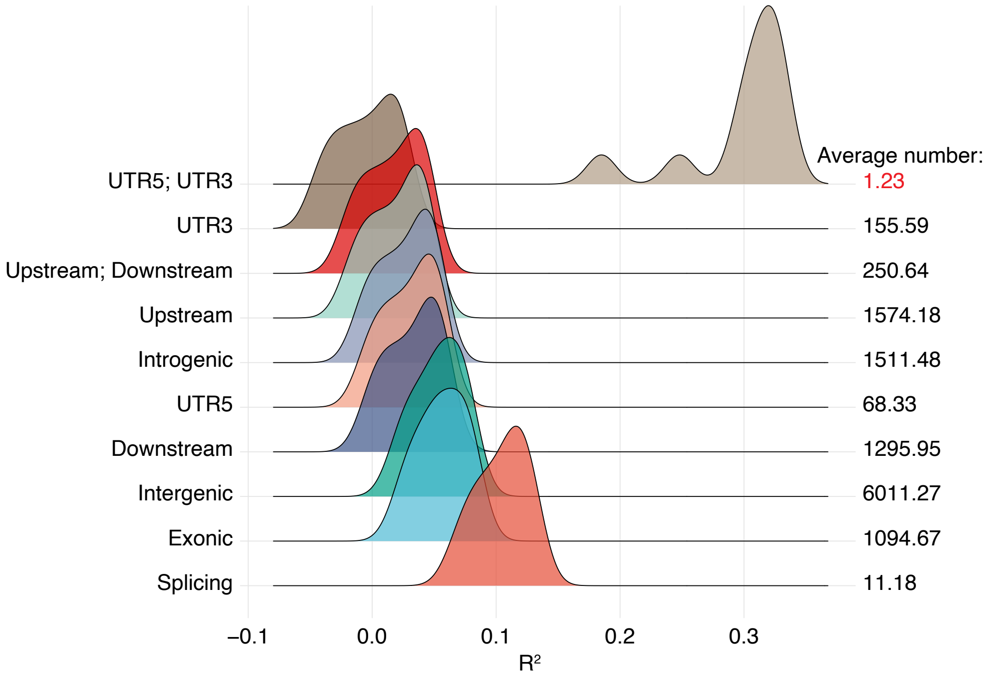


**Supplementary Figure S27.** Adjusted $R^2$ for different types of PVs. The number of parental unique PVs were predicted with additive effect of differentially expressed genes in 12 inbred lines using partial least square model. The PVs were classified into different types based on their location.

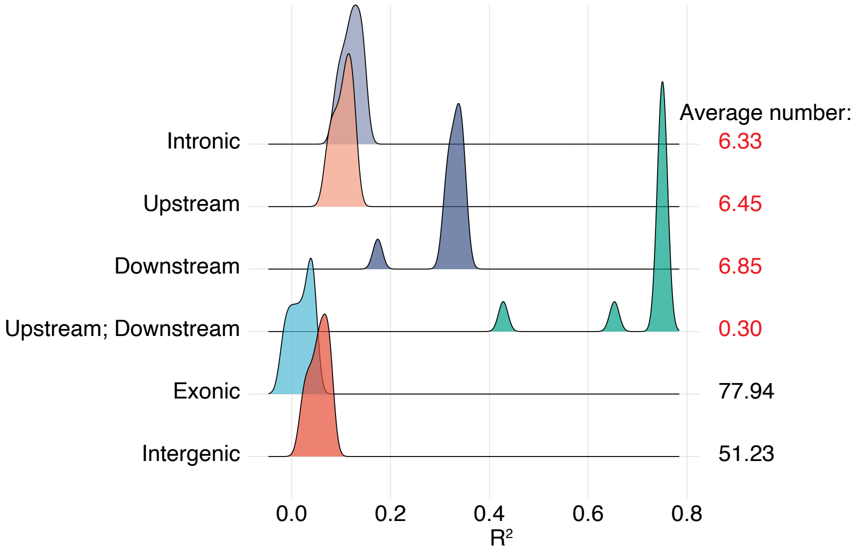


**Supplementary Figure S28.** Adjusted $R^2$ for different types of inversions. The number of parental unique inversions were predicted with additive effect of differentially expressed genes in 12 inbred lines using partial least square model. The inversions were classified into different types based on their location.

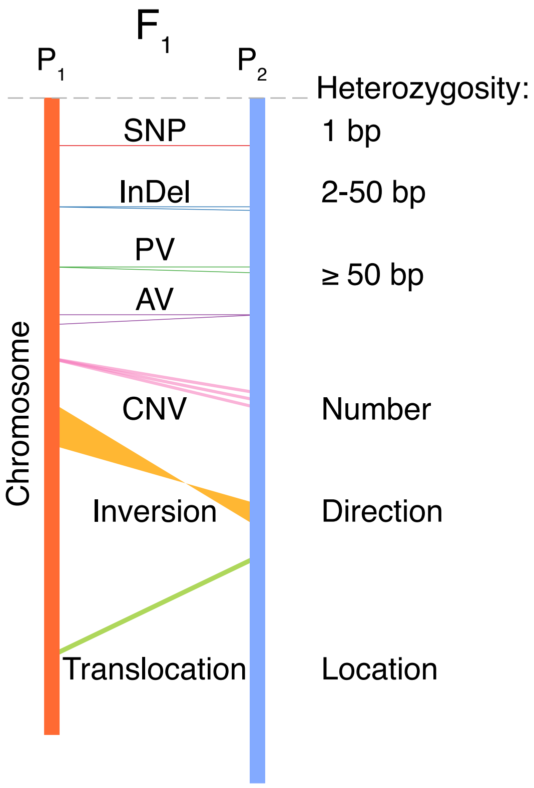


**Supplementary Figure S29.** Heterozygosity of genomic variants in $F_1$ hybrids. Parental genomic variants, including single nucleotide polymorphisms (SNPs), insertions and deletions (InDels), presence/absence variants (PAVs), copy number variants (CNVs), inversions, and translocations, are at heterozygous state in $F_1$ hybrids.
